# Lineage-mixed neurodevelopmental programs in pediatric sarcomas nominate tumor-selective surface antigen pairs

**DOI:** 10.64898/2026.10.28.762504

**Authors:** Alessandro Davini, Xueying Liu, Jackie L. Norrie, Jihye Hwang, Vishwajeeth Pagala, Stephanie D. Byrum, Anthony High, Michael A. Dyer, Heather Sheppard, Anand G. Patel, Paul Geeleher

## Abstract

Pediatric solid tumors have low mutational burden, frequent oncogenic gene fusions, and developmental origins in arrested progenitor cells. Single-cell RNA-seq (scRNA-seq) could represent an ideal tool to assess whether these convergent biological features yield shared therapeutic vulnerabilities. However, while existing tools can extract gene expression programs for pan-tumor single-cell data, interpreting them requires domain expertise across many disciplines and diseases. Here, we analyzed a pan-pediatric scRNA-seq atlas of 266 patient- and xenograft-derived samples across 22 pediatric solid tumor types. To address the bottleneck between gene program identification and interpretation, we developed an AI agent framework built on off-the-shelf large language models (e.g., Gemini 2.5) to interpret gene expression programs at the level of human experts, but at much greater scale and speed. Our agent identified ectopic expression of neuronal marker genes as a unique feature of several fusion-driven sarcomas. This gene expression pattern resulted in tumor-specific co-expression of mixed mesenchymal/neuronal surface markers, providing a plausible therapeutic window to target these tumor types using emerging AND-gated therapeutic strategies, such as bispecific antibody-drug conjugates (ADCs) and bispecific CAR-T cells. As proof of principle, we nominated and validated STEAP1/NPTXR as a Ewing sarcoma candidate cell surface target pair. Together, this work delivers a pan-pediatric solid tumor scRNA-seq atlas, establishes a generalizable and accessible AI-based framework that complements existing single-cell methods by automating interpretation of pan-disease single-cell atlases, and nominates new bispecific ADC/CAR-T targets for catastrophic pediatric cancers.

## INTRODUCTION

Single-cell RNA-sequencing has transformed cancer biology by enabling systematic characterization of cellular heterogeneity and transcriptional programs at unprecedented resolution. Continued improvements in sample processing and computational integration have made it possible to generate atlases comprising millions of cells across hundreds of patients. Computational methods for extracting structure from these data have advanced in parallel. Tools such as matrix factorization and single-cell foundation models (e.g. scGPT, scFoundation) can now extract gene expression programs (GEPs) and cell states from these data^1–4^. However, these methods yield dozens to hundreds of unannotated gene programs, each defined by thousands of gene weights. The task of contextualizing GEPs within our current biological understanding still falls to human experts. This is because extracting biological insight from pan-cancer single-cell atlases requires cross-disease knowledge spanning oncology, immunology, developmental biology, and pharmacology. As atlases grow, we will be faced with a widening mismatch between the volume of available single-cell data and our capacity to convert these data into mechanistic and clinically actionable knowledge.

Large language models (LLMs) offer a potential strategy to narrow this interpretative gap. Recent work has shown that OpenAI’s GPT-4 can annotate cell types from marker gene lists across multiple tissues with accuracy comparable to established tools for gene set analysis^5^, and subsequent benchmarking efforts have evaluated LLMs for cell type and gene set annotation tasks^6^. These studies establish the promise of LLMs as biological annotation tools, but current applications have largely focused on isolated tasks, such as naming clusters or summarizing gene sets. Whether LLMs, used off-the-shelf, can be deployed as a rigorous, scalable interpretation layer for real-world single-cell atlases remains an important open question. In particular, it remains unclear whether LLMs can be deployed to identify recurrent and context-specific gene expression programs (GEPs), to relate these GEPs to tumor biology and clinical metadata, and to nominate experimentally testable therapeutic targets.

Pediatric solid tumors provide a compelling setting in which to develop and evaluate such approaches. Unlike adult carcinomas, which are predominantly epithelial in origin, pediatric solid tumors arise across all three developmental lineages^7,8^. They often have low tumor mutational burden and are frequently driven by oncogenic fusions or mutations that alter transcriptional regulation or chromatin states^9–11^. These features suggest that pediatric cancers may share convergent developmental or transcriptional states, despite originating in distinct tissue contexts. At the same time, their rarity makes cross-tumor analysis especially challenging. Several recent efforts have collected and uniformly processed single-cell datasets from pediatric tumors^12^, yet no detailed analysis of these integrated datasets has been reported.

Here, we performed a detailed analysis of a pan-pediatric solid tumor atlas, comprising single-cell and single-nucleus RNA-seq data generated from 266 St. Jude patient- and patient-derived xenograft (PDX) samples spanning 22 different pediatric solid tumor types. These rare and often difficult-to-obtain biospecimens enabled systematic comparison of malignant, immune, and stromal states across a broad spectrum of pediatric solid tumors. We inferred GEPs using an unsupervised NMF-based framework that embeds malignant and non-malignant compartments into a shared latent space. Rather than training a bespoke interpretation model, we developed a structured prompting framework in which an off-the-shelf LLM (Google’s Gemini 2.5 LLM, state-of-the art at the time of study) acts as an interpretation agent. The agent ingests the output from upstream analyses, including GEP gene loadings, functional enrichments, cell type predictions, malignancy estimates, and clinical metadata. For each GEP, it returns an annotation and a justification to support its label, drawing upon patterns within the pan-tumor dataset and from the literature. Moreover, it synthesized groups of GEP annotations to infer higher-order relationships between GEPs and metadata annotations. Among fusion-driven mesenchymal tumors in our atlas (fusion-positive rhabdomyosarcoma, Ewing sarcoma, CIC-rearranged high-grade sarcomas, and desmoplastic small round cell tumor), our agent identified hybrid mesenchymal-neuronal programs marked by ectopic expression of neuronal genes. These programs expressed tumor-specific co-expression of mesenchymal and neuronal cell-surface markers, suggesting a therapeutic window for emerging AND-gated strategies such as emerging bispecific antibody-drug conjugates and bispecific CAR-T cells. As proof of principle, we nominated and validated STEAP1+NPTXR as a candidate cell-surface target pair in Ewing sarcoma. Together, this work provides an integrated single-cell atlas of pediatric solid tumors, establishes a generalizable AI-agent framework for scalable interpretation of pan-disease single-cell data, and identifies bispecific therapeutic target pairs for catastrophic fusion-driven pediatric sarcomas.

## RESULTS

### The Pediatric Solid Tumor Advanced Repository (PedSTAR)

We analyzed a single-cell/nucleus RNA-sequencing atlas generated from 266 unique pediatric solid tumor samples spanning 22 different tumor types (351 total datasets including replicates; dataset from McEvoy et al, accepted (resource paper, does not include detailed analysis)). This dataset includes common pediatric solid tumors encompassing neuroblastoma (NB), rhabdomyosarcoma (RMS), osteosarcoma (OS), Ewing sarcoma (EWS), retinoblastoma (RB), Wilms tumor (WT), malignant rhabdoid tumor (MRT), as well as multiple rare tumor types, including desmoplastic small round cell tumor (DSRCT) (**Fig. 1a, Supplementary Table 1**). We refer to this dataset as the Pediatric Solid Tumor Advanced Repository (PedSTAR) which, to our knowledge, represents the most comprehensive single-cell atlas of pediatric solid tumors deeply analyzed to date.

**Figure 1.**
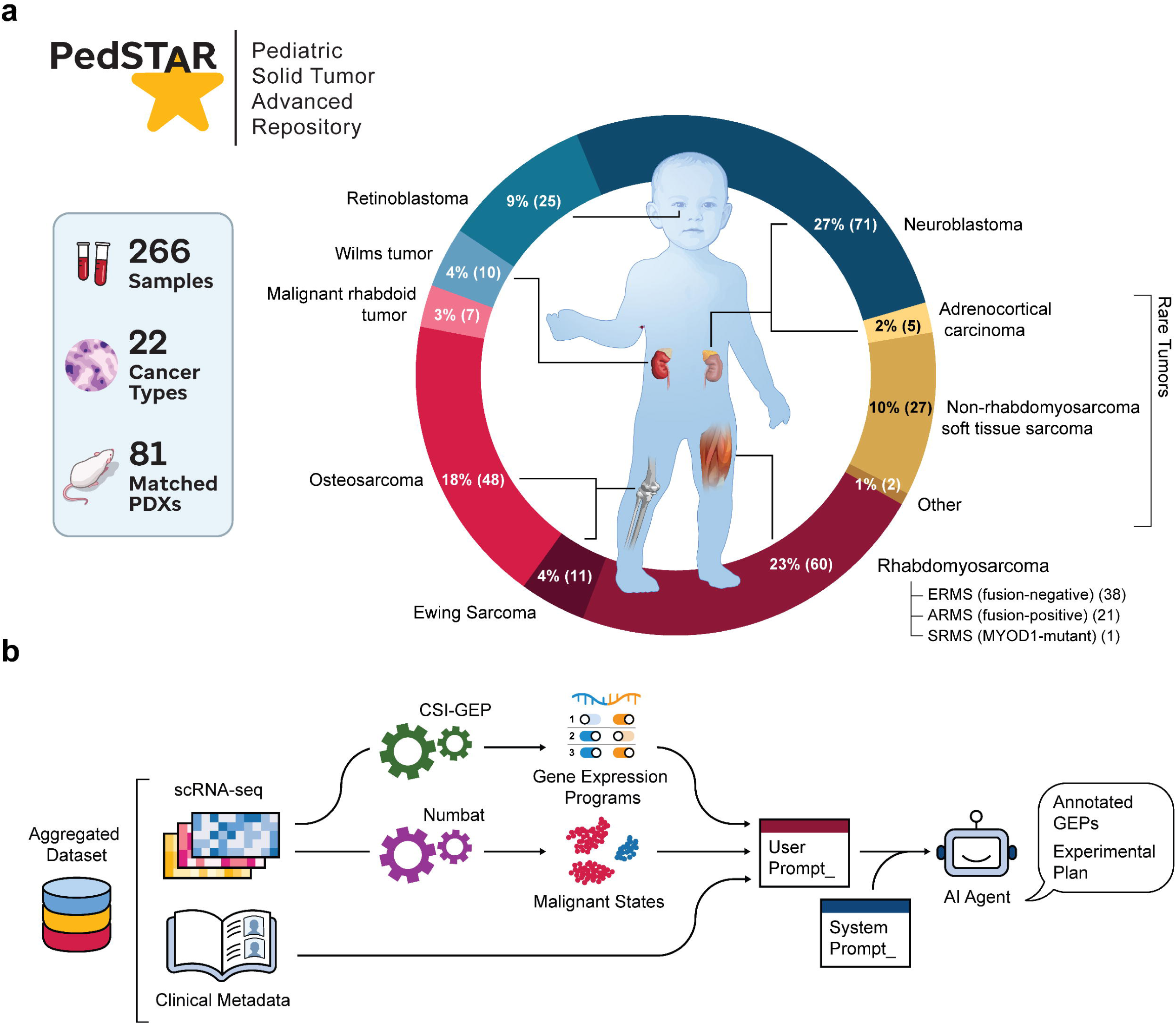
Constructing a pan-pediatric solid tumor single-cell atlas and preparing the dataset for interrogation by our AI agent. **a.** Schematic overview of our PedSTAR dataset, comprising 266 samples across 22 pediatric tumor histologies. **b.** Flowchart summarizing our overall analytical strategy: We used CSI-GEP to recover gene expression programs (GEPs) from the data, Numbat to infer malignant versus non-malignant cells, and collected and assembled all relevant clinical metadata. Our AI agent acts on this information to annotate GEPs and then prioritize downstream biological and experimental hypotheses.

PedSTAR comprises data from 161 unique patient-derived tumor specimens and 105 orthotopic patient-derived xenograft (O-PDX) samples. The atlas integrates data generated from both fresh biospecimens profiled by single-cell RNA-seq (*n* = 109 including replicates) and frozen archival biospecimens profiled by single-nucleus RNA-seq (*n* = 242 including replicates). Notably, 81 patient samples in the atlas have matched O-PDX models represented in the atlas. These O-PDX models are available through the Childhood Solid Tumor Network (CSTN; cstn.stjude.org;^13,14^), providing a practical platform for functional follow-up and experimental validation of hypotheses derived from the atlas.

All samples were processed, quality-controlled, and integrated using a uniform analytical framework, as previously described (McEvoy et al., accepted). The processed data have been made publicly available (see Data Availability). As part of this study we also developed a user-friendly web portal for exploration of summarized PedSTAR data at pscb.stjude.org.

### Defining interpretable gene expression programs in PedSTAR for AI-guided analysis

To enable systematic exploration of the PedSTAR atlas, we first decomposed the sc/snRNA-seq data as a set of interpretable gene expression programs (GEPs). We used Consensus Scalable Inference of Gene Expression Programs (CSI-GEP), a scalable NMF-based package that we previously developed for unsupervised discovery of recurrent transcriptional programs in large single-cell datasets^1,4^. In this setting, individual GEPs can represent discrete cell types as well as coordinated transcriptional features shared *across* cell types, such as pathway or gene set activity.

CSI-GEP identified 134 GEPs across PedSTAR. To assign GEP activity to each single cell, we fit Gaussian Mixture Models (GMMs) to the activity score distribution of each GEP to classify each cell as GEP-expressing or -nonexpressing (**Supplementary Figure 1**). We then inferred malignant versus non-malignant state of each cell using Numbat^15^, allowing us to distinguish GEPs predominantly in malignant cells from those GEPs arising in immune, stromal, or other non-malignant compartments. Finally, we harmonized sample-level clinical and pathological metadata, including tumor type, biospecimen source, treatment status, metastatic site, and outcome (**Supplementary Table 1**). Altogether, the GEP definitions (gene identities and weights), cell-level GEP assignments, malignant/non-malignant classification, and harmonized clinical data formed the structured input for the AI agent’s reasoning engine (**Fig. 1b**). This representation enabled the agent to interrogate the atlas at the level of transcriptional programs, rather than individual genes, supporting scalable identification of convergent expression programs, disease-specific programs, and clinically relevant associations.

### Developing a context-aware AI agent for GEP annotation

To annotate and interpret the 134 GEPs identified within PedSTAR, we developed a structured prompting strategy for the AI agent. The strategy combined a ‘system prompt’, which defined the overall task, input schema, output format, and available metadata, with a ‘user prompt’, which were generated for each GEP program (**Fig. 2a; Supplementary Files 1 and 2**). Each user prompt supplied the biological information needed to interpret an individual GEP.

**Figure 2.**
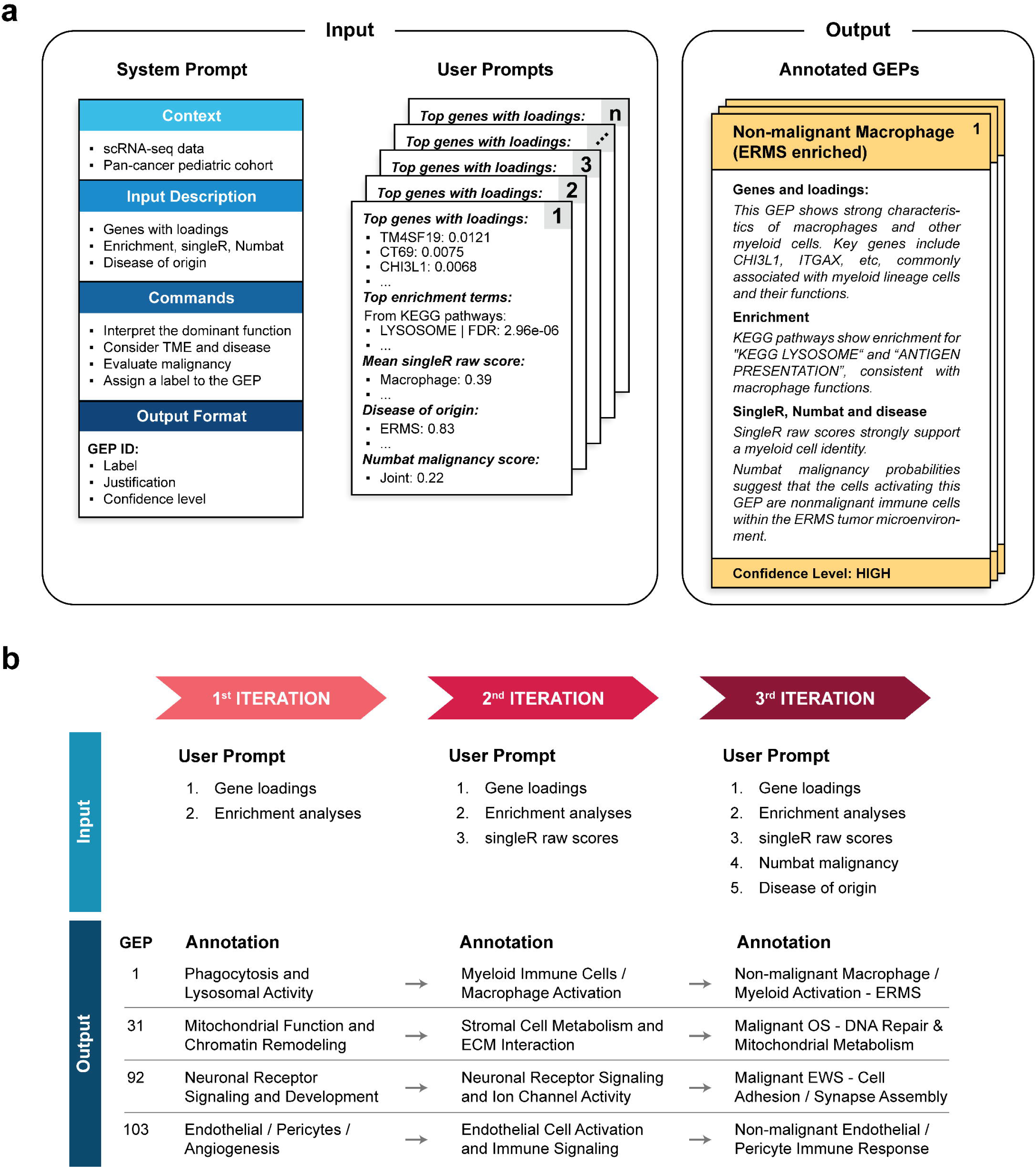
Developing an AI agent for annotating the PedSTAR GEPs. **a.** Schematic illustrating the structure of the system and user prompts used to guide the AI agent. The broad system prompt defined the analysis context and desired output format, while the GEP-specific user prompt provided top ranked genes by NMF loading, gene set enrichment results, SingleR cell-type prediction scores, Numbat malignancy probabilities, and disease-of-origin information for each GEP’s top-activating cells. The AI agent integrated these inputs to assign a label, justification, and confidence level for each GEP. **b.** We evaluated three independent prompt configurations that provided progressively richer biological context. In the 1^st^ iteration, the model received only gene loadings and gene set functional enrichment results, producing coarse functional assignments. In the 2^nd^ iteration, we added SingleR scores, refining cell-type specificity. In the 3^rd^ iteration, we further included malignancy probabilities and disease-of-origin information, enabling clearer separation of malignant and non-malignant programs and highlighting detailed tumor-specific features.

For each GEP, the user prompt included the top-ranked gene loadings from CSI-GEP’s NMF output, corresponding to the program-defining overexpressed marker genes, as well as significant functional enrichments from multiple gene set databases, including Gene Ontology and KEGG (**Fig. 2a**). We also incorporate orthogonal annotations derived from single-cell data, including SingleR cell-type prediction scores^16^, Numbat malignancy probabilities^15^, and the tumor type for cells classified as GEP-expressing by GMM-based thresholding. Using these inputs, the agent was instructed to infer the primary biological function of each GEP, assign a concise descriptive label, provide a rationale for the annotation, and report a confidence rating.

Next, we evaluated how the scope of biological context supplied to the agent impacted the resulting annotations. We tested three user prompt configurations that provided iteratively richer information. In the first iteration, user prompts included only gene loadings and functional enrichment terms. This iteration produced coarse annotations that largely reflected dominant pathway or process-level signals within each GEP (**Fig. 2b**). In the second iteration, we added SingleR-based cell-type prediction scores, which shifted the annotations towards more cell type-centered labels (**Fig. 2b**). In the third iteration, we further added Numbat-derived malignancy probabilities and disease-of-origin information for GEP-positive cells. This most context-rich configuration produced the most informative annotations, allowing the agent to better distinguish between malignant and non-malignant programs and to recognize tumor-type-specific patterns. Overall, increasing the amount of structured biological context supplied through the user prompt improved the specificity and interpretability of the agent’s annotations (**Fig. 2b; Supplementary Table 2**). These improvements were quantified in benchmarking analyses described in the next section.

### Benchmarking the AI agent’s GEP annotations on a published neuroblastoma dataset previously annotated by human subject matter experts

Before applying the AI agent to the full PedSTAR atlas, we benchmarked its GEP annotations using a smaller, previously published scRNA-seq dataset with expert human annotations. For this analysis, we selected the GOSH neuroblastoma dataset^17^, which includes five samples and 41 GEPs previously identified using an NMF-based methodology similar to CSI-GEP^1^. Because these GEPs were annotated manually by two blinded subject experts before the widespread availability of LLMs, they provide an independent reference set for evaluating AI-generated annotations.

We first tested whether current LLMs could recover human assigned labels in a constrained label-matching task. We evaluated both Google’s Gemini 2.5 and OpenAI’s GPT-5, two best-in-class models available programmatically through an API at the time of this study. For each GEP, we supplied each LLM with structured input files containing the same prompt-derived information described above, together with a shuffled list of the 41 published human-assigned labels. The models were asked to select the best matching label for each GEP. We repeated this evaluation 100 times for each model to assess performance across repeated prompt executions (**Fig. 3a; Supplementary Figure 2**). In this task, Gemini 2.5 substantially outperformed GPT-5, assigning the published label correctly for 60% of GEPs, compared to ∼2% for GPT-5 (**Fig. 3a**). Notably, GEPs that human experts had originally rated as “low-confidence” were also among the programs most frequently misassigned by the models (**Supplementary Figure 2**), suggesting that ambiguous programs posed similar challenges for human and AI-based annotation.

**Figure 3.**
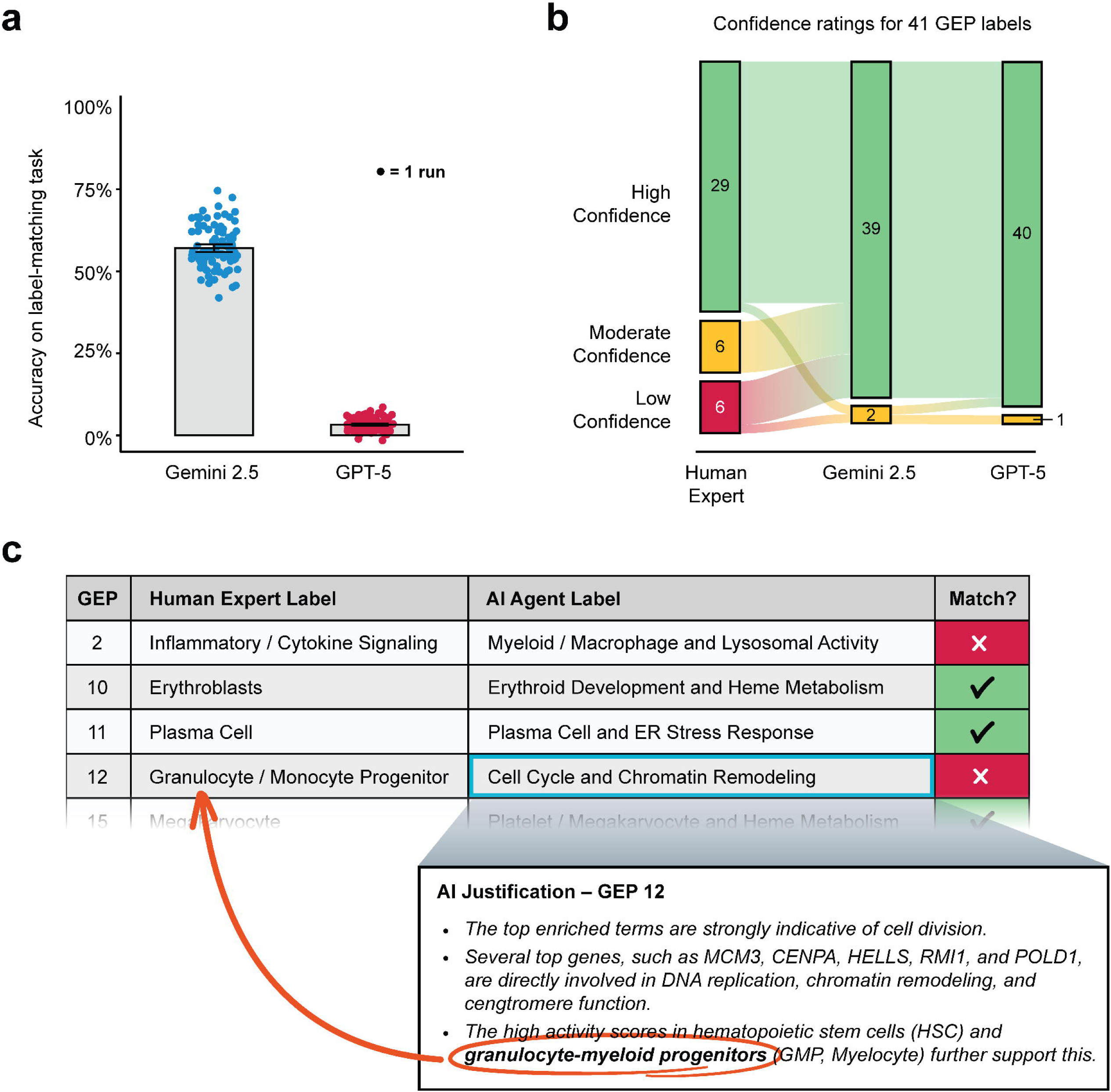
Benchmarking the AI agent’s GEP annotations on a published neuroblastoma dataset that was previously annotated by human subject matter experts. **a.** Barplot comparing accuracy across 100 attempts by Gemini and GPT-5 agents to select the identical previous human-assigned GEP label in GOSH. **b.** Sankey bar chart showing confidence scores assigned by human annotators, GPT-5, and Gemini during the *de novo* (qualitative) GEP annotation task. **c.** Tables summarizing AI-human concordance. Left panel: examples of qualitatively matching GEP label annotations. Right panel: examples of discordant annotations, with AI justifications revealing reliance on similar biological evidence but emphasizing e.g. functional processes over cell-type specificity.

We next evaluated a more realistic de novo annotation task, in which the models generated GEP labels directly from the structured biological inputs rather than selecting from a predefined list. For each GEP, the models were asked to generate a concise label, justify the annotation, and provide a confidence rating. Although Gemini 2.5 outperformed GPT-5 in the previous constrained label-matching task, the two LLMs performed similarly in this *de novo* task (**Supplementary Table 3**). Both models rated nearly all of their annotations as “high confidence” (**Fig. 3b**). Manual review showed that the AI-generated labels were broadly concordant with the published expert labels, with major discrepancies observed for only 6 of 41 GEPs.

For those discordant program labels, differences between human and AI annotations generally reflected naming conventions rather than fundamentally different interpretations of the underlying data. In several cases, the AI-generated rationale identified the same cell population or biological signal emphasized by the human label but assigned a more function-oriented name. For example, human experts labeled GEP 12, “Granulocyte monocyte progenitor,” whereas the AI models assigned the label “Cell cycle and chromatin remodeling”. Despite the divergent label, the AI justification cited “high activity…in granulocyte-myeloid progenitors,” indicating that the model recognized the relevant cellular context while emphasizing the dominant cell cycle and chromatin-associated features of the program (**Fig. 3c**). Similarly, GEP 10 was labeled “Erythroblasts” by human experts, whereas the AI labeled the same program “Erythroid Development and Heme Metabolism” (**Fig. 3c; Supplementary Table 3**). In these cases, the human labels tended to emphasize cell type identity, while the AI emphasized biological process or function (**Fig. 3c**).

Collectively, these benchmarks show that the agent can generate GEP annotations that are concordant with human experts, while also exposing systematic differences in annotation style. These results supported deployment of the agent for PedSTAR, where the scale and diversity of the atlas make comprehensive manual interpretation a major bottleneck.

### PedSTAR-wide annotation identifies convergent neurodevelopmental programs in fusion-driven sarcomas

Encouraged by the agent’s performance in the benchmarking tasks, we next applied it to the full PedSTAR atlas, annotating all 134 GEPs (**Supplementary Table 4**). The agent’s annotations, together with the underlying inputs used to generate them, are available for exploration at pscb.stjude.org. The agent assigned high confidence to 129 of 134 annotations (96.3%), with four programs assigned moderate confidence and one assigned low confidence (**Fig. 4a**). The only low confidence program, GEP 132, was expressed across cells from 15 tumor types and lacked significant gene set enrichments. A similar pattern was observed among GEPs assigned moderate confidence, such as GEP 130, which also showed broad cross-tumor activity without clear functional enrichment. Thus, the agent’s confidence scores aligned with the amount of biological information available to interpret each program.

**Figure 4.**
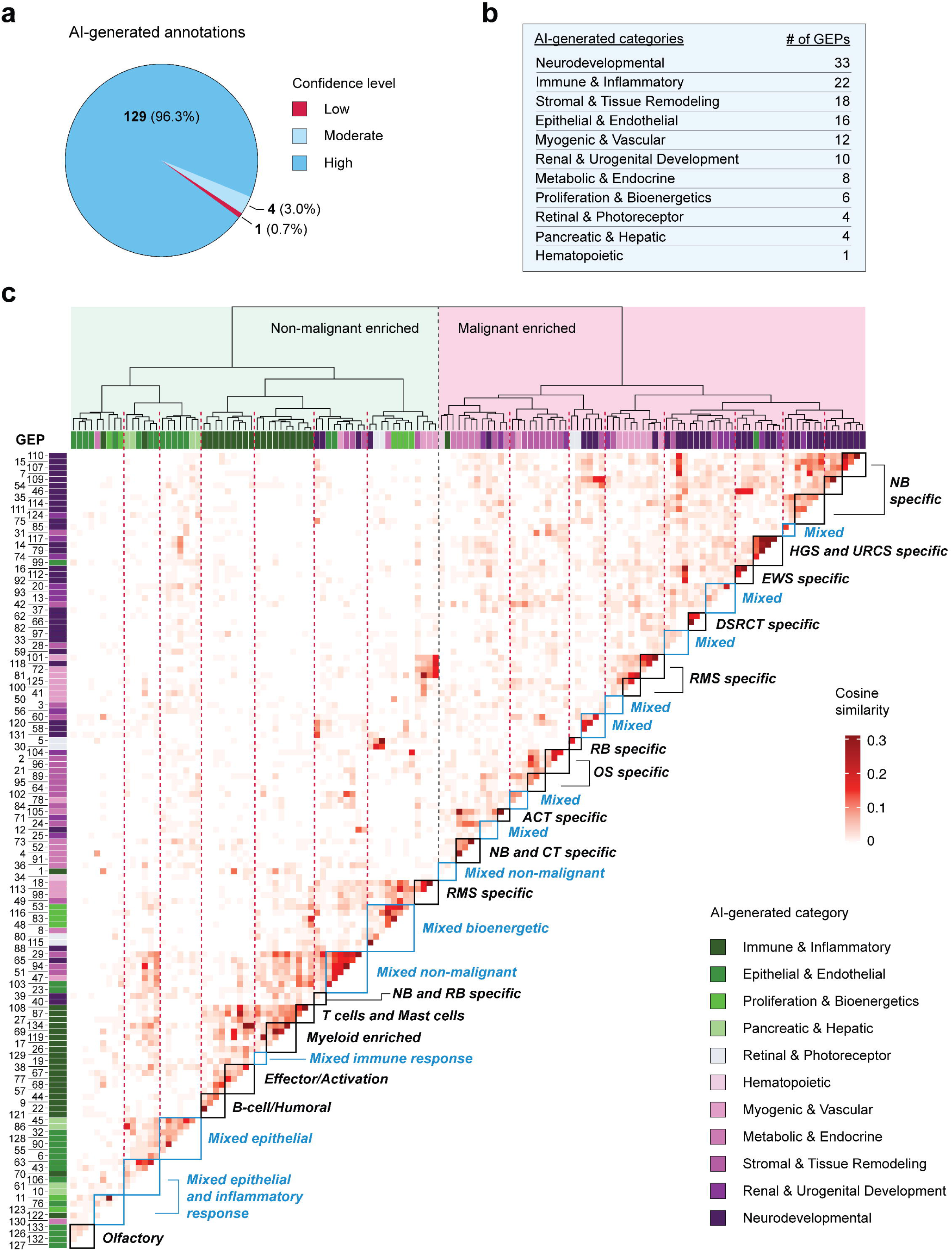
Atlas-wide annotation, categorization, and interpretation of 134 GEPs by the AI agent. **a.** Pie chart showing distribution of AI’s subjective confidence levels for all annotated GEPs. **b.** Table showing an overview of the 11 biological categories identified by the AI agent across the 134 GEPs, including neurodevelopmental, immune and inflammatory, stromal and tissue remodeling, epithelial and endothelial, myogenic and vascular, renal and urogenital development, metabolic and endocrine, proliferation and bioenergetics, retinal and photoreceptor, pancreatic and hepatic, and hematopoietic programs. **c.** Heatmap showing the cosine similarity (colors) generated from the top 50 gene loading scores defining each GEP, grouped using unsupervised clustering (dendrogram).

To organize the high confidence annotations, we also prompted the agent to group GEPs according to their dominant biological features. This yielded 11 broad categories (**Fig. 4b**; pscb.stjude.org), including 22 “Immune and inflammatory” GEPs, consistent with the expected cellular composition of the tumor microenvironment. To further examine global relationships among programs, we computed cosine similarity for all pairwise combinations of GEPs using their top 50 gene loading scores (8,911 pairs total) and then applied hierarchical clustering. This analysis revealed a clear separation between malignant and non-malignant programs (**Fig. 4c**).

Non-malignant GEPs clustered into expected tumor microenvironment compartments, including immune, epithelial, and stromal programs. However, shared features and co-expression patterns became apparent within both malignant and non-malignant program groups, suggesting that meaningful higher-order relationships existed beyond primary annotations.

Although the agent generated accurate labels and descriptions for individual GEPs, interpreting thousands of pairwise relationships and placing these patterns into biological context remained a major analytical challenge. We therefore performed a second atlas-level interpretation step in which the agent reasoned over the full set of GEP labels, textual justifications, co-expression patterns, and associated metadata. The agent was instructed to connect these observations to existing literature, and to summarize major biological themes and experimental opportunities emerging from the data (**Supplementary File 3**). In this meta-annotation analysis, the agent highlighted several broad patterns, including tightly co-regulated GEPs defining the malignant neural crest identity of neuroblastoma cells and recurrent oxidative phosphorylation programs in neuroblastoma and retinoblastoma, suggesting potential metabolic dependencies that may be therapeutically exploitable.

We became particularly interested in the agent’s assignment of 33 GEPs to a neurodevelopmental category. As expected, this group included neuroblastoma and retinoblastoma programs. Unexpectedly, however, it also included malignant cells from multiple pediatric sarcomas, which are classically considered mesenchymal tumors. Across both the primary GEP annotations and atlas-wide meta-annotation, the agent highlighted neuron-like or neurodevelopmental features in Ewing sarcoma (EWS; GEPs 16, 46, 92, 97, and 112), desmoplastic small round cell tumor (DSRCT; GEPs 62, 66, and 82), CIC-rearranged high-grade sarcomas (HGS; GEPs 14 and 79), and alveolar rhabdomyosarcoma (ARMS; GEP 118) (**Supplementary Figure 3**).

Human review confirmed that these neurodevelopmental GEP assignments were supported by the underlying gene loadings and enrichment results. For example, among the top mSigDB cell-type gene set enrichments for GEP 16 in Ewing sarcoma and GEP 62 in DSRCT were neuronal development programs derived from the La Manno et al. single-cell brain atlas^18^. GEP 16 was enriched for “Midbrain neurotypes serotonergic” (odds ratio = 25.49, *FWER* = 6.98 × 10^-10^), whereas GEP 62 was enriched for “Midbrain neurotypes GABAergic” (odds ratio = 11.56, *FWER* = 2.4 × 10^-4^). Once flagged by the agent, these enrichments were verified by human inspection and can be corroborated through our PedSTAR web portal (pscb.stjude.org).

Notably, the sarcoma-associated neurodevelopmental GEPs arose broadly across fusion-driven sarcomas (**Supplementary Table 5**), including tumors characterized by the oncogenic transcription fusions *EWS-FLI1* in Ewing sarcoma, *EWS-WT1 in* DSRCT, *PAX3/7-FOXO1* in ARMS, and *CIC-DUX4 in* high-grade sarcoma (**Supplementary Table 1**). Importantly, these findings are consistent with prior literature reporting neuronal-like gene expression individually in Ewing sarcoma^19–21^, DSRCT^22–24^, and alveolar rhabdomyosarcoma^25–27^. However, ectopic expression of neuronal marker genes as a shared feature of fusion-driven pediatric sarcomas has not been systematically described nor has a therapeutic strategy been proposed to exploit this biology.

### Dissecting neuron-relevant gene expression in fusion positive pediatric sarcomas

We next asked whether the aberrant neuronal features observed across fusion-positive pediatric sarcomas reflected a shared neurodevelopmental program or instead represented disease-specific patterns of ectopic neuronal gene expression. Distinguishing between these possibilities could provide insight into the mechanistic basis of these programs and clarify whether they might be therapeutically exploitable.

We first compiled a broad neuron-associated gene set by combining genes annotated to the Gene Ontology term “neuron differentiation” with synaptic genes curated in the SynGO database^28^ (**Supplementary Table 6**; *n* = 2,497 genes). We assessed the loading scores for these genes across the 11 neurodevelopmental GEPs identified in fusion-driven sarcomas. This analysis centered on the 110 neuron-associated genes that ranked in the top 50 markers for at least one of the 11 GEPs (**Fig. 5a**). These ectopically expressed neuronal marker genes were not limited to specific programs or individual patients. Instead, they recurred across multiple samples within each disease, suggesting that neuronal gene expression is a reproducible feature of these sarcomas. However, the specific neuronal markers varied across Ewing sarcoma, DSRCT, ARMS, and CIC-rearranged sarcomas.

**Figure 5.**
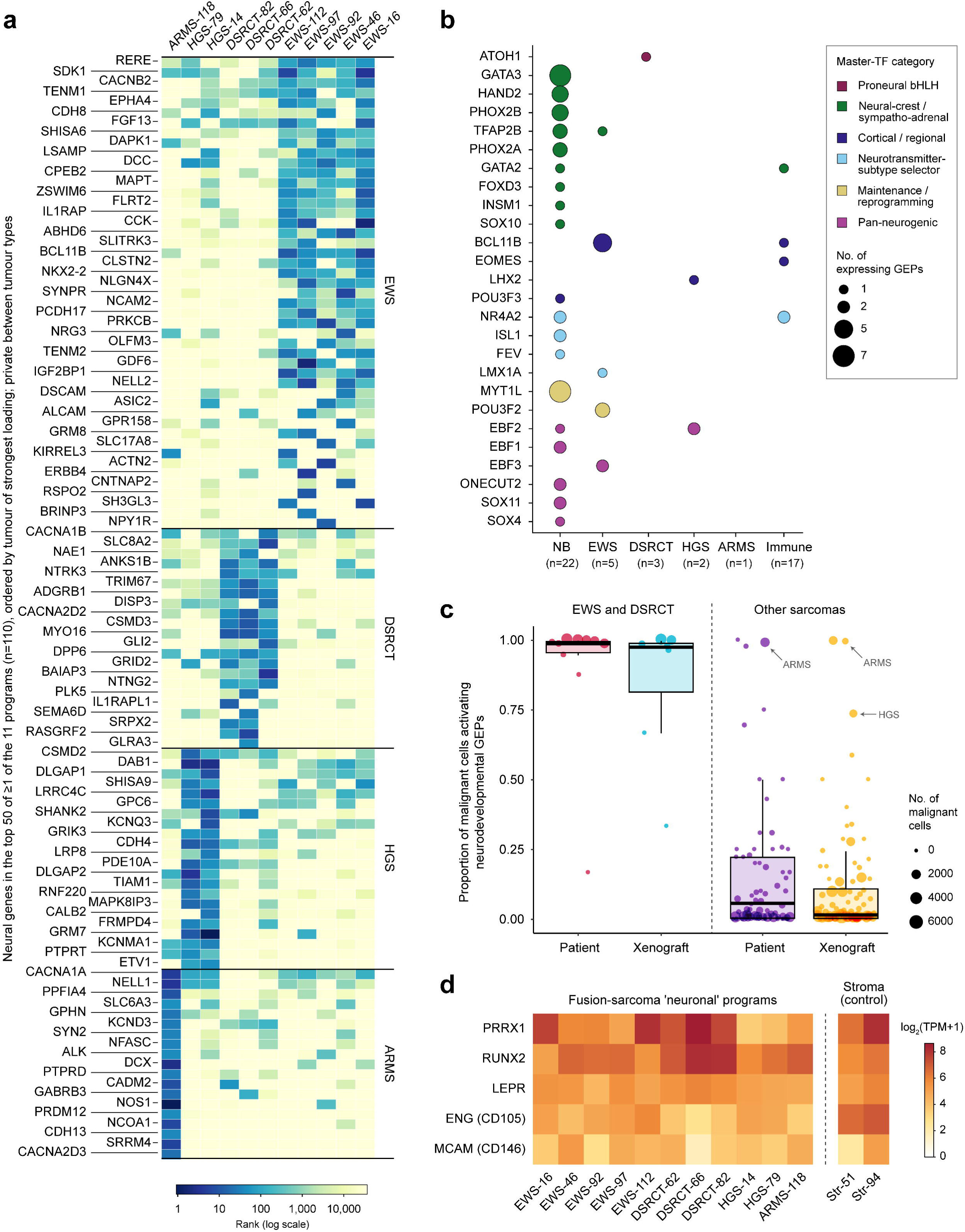
Dissecting ectopic neuronal gene expression in malignant sarcoma cells. **a.** Heatmap showing the per-GEP loading rank of neuron-associated genes (columns) across the 11 fusion-driven sarcoma neurodevelopmental GEPs (rows). Color encodes each gene’s loading rank within a GEP on a log_10_ scale. Genes are ordered by the tumor type in which they are most strongly loaded and then by mean loading rank across GEPs. **b.** Dot plot summarizing how frequently curated neurodevelopmental master TFs (rows) appeared among the top 20 TFs by loading score in each GEP group (columns). Dot size is proportional to the number of GEPs in the group containing that factor. Dot color denotes the factor’s developmental subcategory. **c.** Boxplot showing the proportion of malignant cells activating neurodevelopmental programs in Ewing and DSRCT samples (left) and other sarcomas (right). **d.** Heatmap of gene expression per-GEP TPM values for five classical mesenchymal markers not expected to be neural-confounded, across the 11 fusion-driven sarcoma neurodevelopmental GEPs (left block) and two non-malignant mesenchymal-stromal/fibroblast GEPs (stroma control; GEPs 51 and 94; right block). Columns are individual GEPs; color encodes log_2_ transcripts per million (TPM) values.

To determine whether these patterns reflected activation of canonical neurodevelopmental regulatory circuitry, we manually curated a list of 58 neurodevelopmentally relevant master transcription factors (TFs) from the literature. These included proneural bHLH fate determinants, neural crest and autonomic regulators, cortical patterning and projection-neuron specification factors, neuronal subtype and terminal differentiation regulators, and neuronal identity or reprogramming factors (**Supplementary Table 7**). We assessed how frequently these TFs appeared among the top 20 TFs by loading scores in the fusion-driven sarcoma neurodevelopmental GEPs. As a positive comparator, neuroblastoma programs reproducibly assembled a coherent neural-crest/sympathoadrenal TF module, marked by expression of *HAND2*, *PHOX2B*, *GATA3*, and related regulators. In contrast, neuronal TF expression in fusion-driven sarcomas appeared idiosyncratic and did not converge on a consistent regulatory module that resembled normal neuronal differentiation (**Fig. 5b**).

These findings suggest that ectopic expression of neuronal genes in fusion-driven sarcomas may arise from transcriptional dysregulation imposed by oncogenic fusion drivers rather than from activation of canonical neurodevelopmental differentiation program. This interpretation is consistent with previous reports in Ewing sarcoma showing that the neuronal transcriptional repressor *BCL11B*, which was highly ranked across all Ewing sarcoma neurodevelopmental GEPs in our analysis, is a direct regulatory target of the *EWS-FLI1* fusion gene^29,30^. More broadly, our data suggest that fusion oncogenes induce disease-specific neuronal gene expression patterns that are distinct from physiologic developmental programs.

We next asked whether neurodevelopmental programs were expressed broadly across malignant cells within individual tumors, since widespread expression would enhance the therapeutic relevance of these programs. For each sarcoma sample, we estimated the proportion of Numbat-classified malignant cells that expressed one of the 33 neurodevelopmental GEPs (**Supplementary Table 5**). Nearly all malignant cells in Ewing sarcoma and DSRCT expressed a neurodevelopmental program, while the estimated proportion exceeded 75% in ARMS and CIC-rearranged high-grade sarcoma (**Fig. 5c**; **Supplementary Table 5**). This analysis also revealed little difference between patient tumors and matched O-PDX models in the prevalence of neurodevelopmental program expression. Thus, ectopic neuronal gene expression appears to be maintained after orthotopic transplantation into mice, supporting the use of O-PDXs as model systems for functional studies of these aberrant transcriptional states.

Despite acquiring neuronal features, fusion-driven sarcoma cells retained transcriptional hallmarks of their mesenchymal origins. Classical mesenchymal markers including *PRRX1*^31^, *RUNX2*^32^, *LEPR*^33^, *ENG* [CD105;^34^], *MCAM* [CD146;^35^], were expressed in cells assigned to the 11 fusion-driven sarcoma GEPs at levels comparable to those observed in stromal and fibroblast GEP-expressing cells (**Fig. 5d**). Collectively, these data indicate that fusion-driven pediatric sarcomas can occupy mixed mesenchymal-neuronal transcriptional states.

### Neuronal and mesenchymal candidate target pairs are aberrantly co-expressed on the surface of EWS cells

These lineage-mixed states may create therapeutic opportunities. In normal tissues, mesenchymal and neuronal surface proteins are lineage-segregated. In fusion-driven sarcomas, however, these markers may be aberrantly co-expressed on the same malignant cells, creating a potential therapeutic window for emerging AND-gated strategies such as bispecific CAR-T cells and bispecific antibody-drug conjugates. We reasoned that gene loading scores from the neurodevelopmental GEPs could be used to prioritize overexpressed cell-surface antigen pairs with potential therapeutic relevance. We focused first on Ewing sarcoma, where ectopic neuronal gene expression has been extensively described. We aggregated the top 100 genes by loading score from each of the five Ewing sarcoma neurodevelopmental GEPs and filtered them for predicted plasma membrane localization using curated annotations from UniProt, Gene Ontology, and the COMPARTMENTS subcellular localization database^36–39^. We then annotated candidate surface genes according to their reported biological roles and tissue expression patterns, distinguishing neuronal lineage-associated genes from canonical EWS-or mesenchymal-associated surface genes. To identify surface markers that were co-expressed in EWS cells but segregated across normal tissues, which would suggest a potential therapeutic window, we examined baseline expression of the candidate surface genes across normal human tissues using GTEx RNA-seq data^40^. This analysis prioritized six candidate neuronal surface genes with tissue-restricted expression in GTEx normal brain: *NPTXR*, *MLC1*, *NLGN1*, *TMEM178B*, *CACNA1B*, and *TENM1.* Similarly, we identified two mesenchymal surface genes that were predominantly expressed in mesenchymal tissues and minimally expressed in normal brain: *STEAP1* and *SDK1* (**Fig. 6a**).

**Figure 6.**
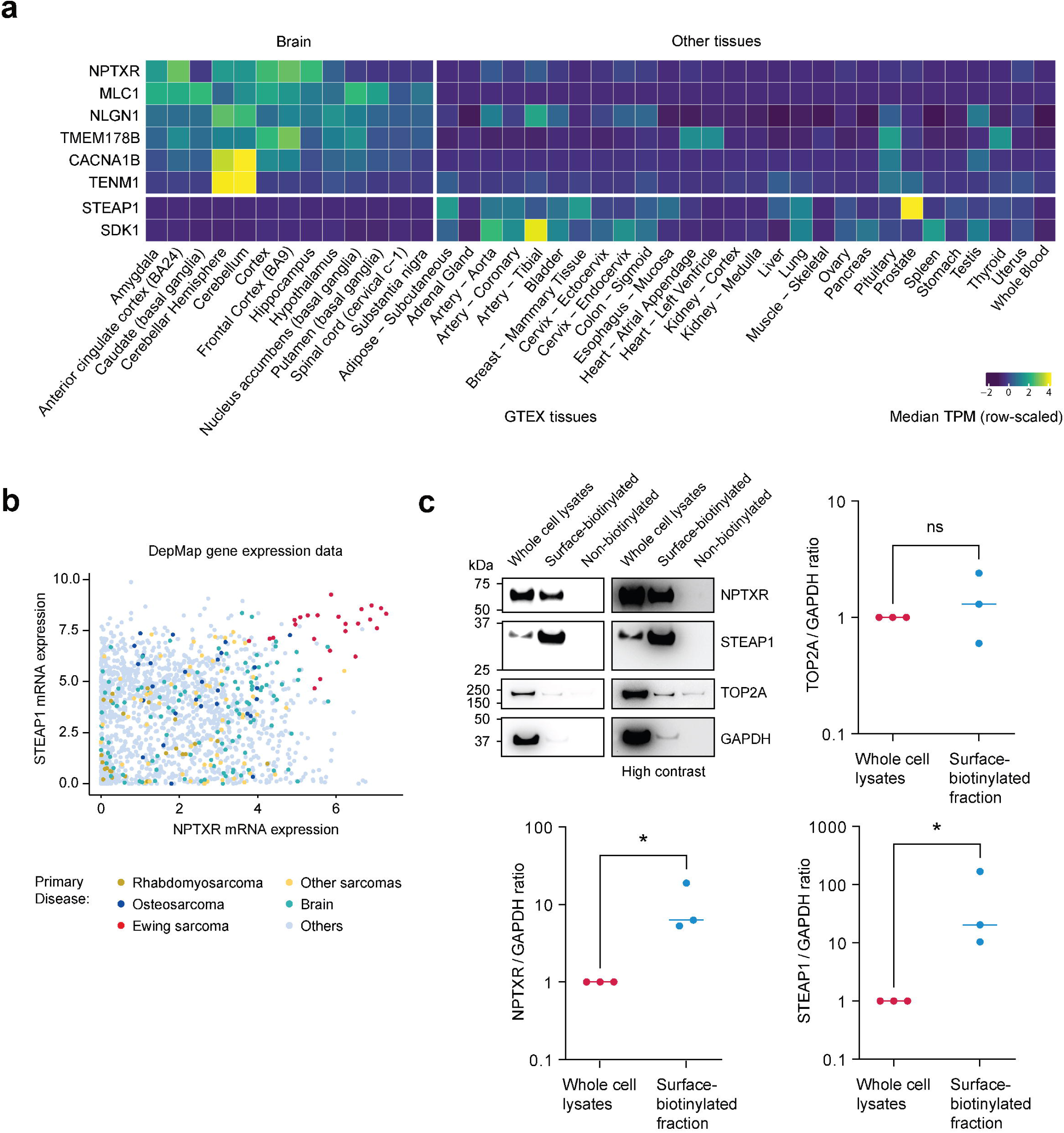
Co-expression of neuronal and Ewing sarcoma programs identifies bispecific target candidates. **a.** Expression of candidate surface markers across GTEx tissues (row-scaled median TPM). *NPTXR*, *MLC1*, *NLGN1*, *TMEM178B*, *CACNA1B*, and *TENM1* show predominant expression in brain tissues, while *STEAP1* and *SDK1* show minimal expression in brain. **b.** DepMap gene expression data illustrating co-overexpression of *STEAP1* and *NPTXR* in cell lines. **c.** Representative western blots of whole-cell lysates, surface-biotinylated fractions, and matched non-biotinylated controls for NPTXR, STEAP1, TOP2A (negative control), and GAPDH (top left panel). High-contrast images are shown to facilitate visualization of residual signal. The remaining panels show densitometric quantification (y axis) of NPTXR, STEAP1, and TOP2A normalized to GAPDH. Each point represents a biologically independent experiment, with whole-cell lysate values normalized to 1 for each experiment and horizontal lines indicate the mean. * indicates *P* < 0.05 from paired one-tailed t-test; *ns*, not significant.

From this candidate gene list, we selected *STEAP1* and *NPTXR* as the mixed-lineage surface antigen pair with the strongest immediate translational rationale. *STEAP1* is a known Ewing sarcoma-associated surface antigen and has been explored as a target for CAR-T approaches^41,42^. NPTXR, a neuronal pentraxin receptor^43^, is beginning to be evaluated as an ADC target in gastrointestinal and other cancers but has not yet been explored in Ewing sarcoma. To test whether this atlas-defined candidate pair was preserved in biologically independent Ewing sarcoma models, we examined bulk transcriptomic data from cancer cell lines, reported in the DepMap database^44^. EWS cell lines showed a striking and selective co-overexpression of *STEAP1* and *NPTXR* (**Fig. 6b**). Consistent with this observation, DepMap protein expression data also supported co-expression of *STEAP1* and *NPTXR*, with the strongest signal observed in a EWS cell line (**Supplementary Figure 4**). Furthermore, we analyzed the St. Jude PeCan bulk RNA-seq dataset^45^ and found that (*n* = 35) Ewing sarcoma patient tumors strongly overexpressed both *NPTXR* and *STEAP1* relative to (*n* = 689) other pediatric solid tumors (**Supplementary Figure 5**). Together with our analysis of PedSTAR dataset, these independent datasets suggest that aberrant co-expression of neuronal and mesenchymal surface antigens is a stable feature of EWS, maintained even in long-established *in vitro* cell line models. Finally, we performed cell-surface biotinylation followed by NeutrAvidin enrichment and immunoblotting, confirming expression of NPTXR in the surface fraction across three independent biological replicates (TC32 Ewing sarcoma cells). As a positive control, STEAP1 was enriched, whereas intracellular proteins GAPDH and TOP2A were depleted (**Fig. 6c**). Data-independent acquisition (DIA) mass spectrometry of two independent surface-enriched preparations identified STEAP1 in both experiments, along with numerous known/expected surface proteins (**Supplementary Tables 8 and 9**).

To further evaluate STEAP1/NPTXR as a candidate bispecific target pair in Ewing sarcoma, we next performed immunohistochemical staining of tissue collected from O-PDXs and patient samples. We first examined O-PDXs generated from Ewing sarcoma, rhabdomyosarcoma, neuroblastoma, and osteosarcoma tumors (**Supplementary Figure 6a**). While we identified examples of NPTXR-positivity in neuroblastoma and rhabdomyosarcoma and STEAP1-positivity in DSRCT, only Ewing sarcoma O-PDXs were consistently dual-positive for both markers. We performed dual-immunofluorescence staining of EWS O-PDXs and noted striking colocalization along with a cell surface pattern of staining (**Fig. 7a and Supplementary Figure 6b**). Next, we co-stained tissue microarrays collected from patient biospecimens of Ewing sarcoma (n = 20), DSRCT (n = 11), Wilms tumor (n = 13), osteosarcoma (n = 15), neuroblastoma (n = 18), and rhabdomyosarcoma (n = 21) (**Fig. 7b and Supplementary Table 10**). Notably, we observed dual-positive STEAP1/NPTXR staining in EWS (14/20; 70%) and rare DSRCT samples (2/11; 18.2%) **(Fig. 7c**). Together, these analyses nominate STEAP1/NPTXR as a validated EWS surface antigen pair with potential relevance for bispecific ADC or CAR-T development. More broadly, this proof-of-principle illustrates how atlas-scale single-cell transcriptomic data and AI-guided GEP interpretation can identify tumor-selective antigen combinations emerging from aberrant mixed-lineage transcriptional states.

**Figure 7.**
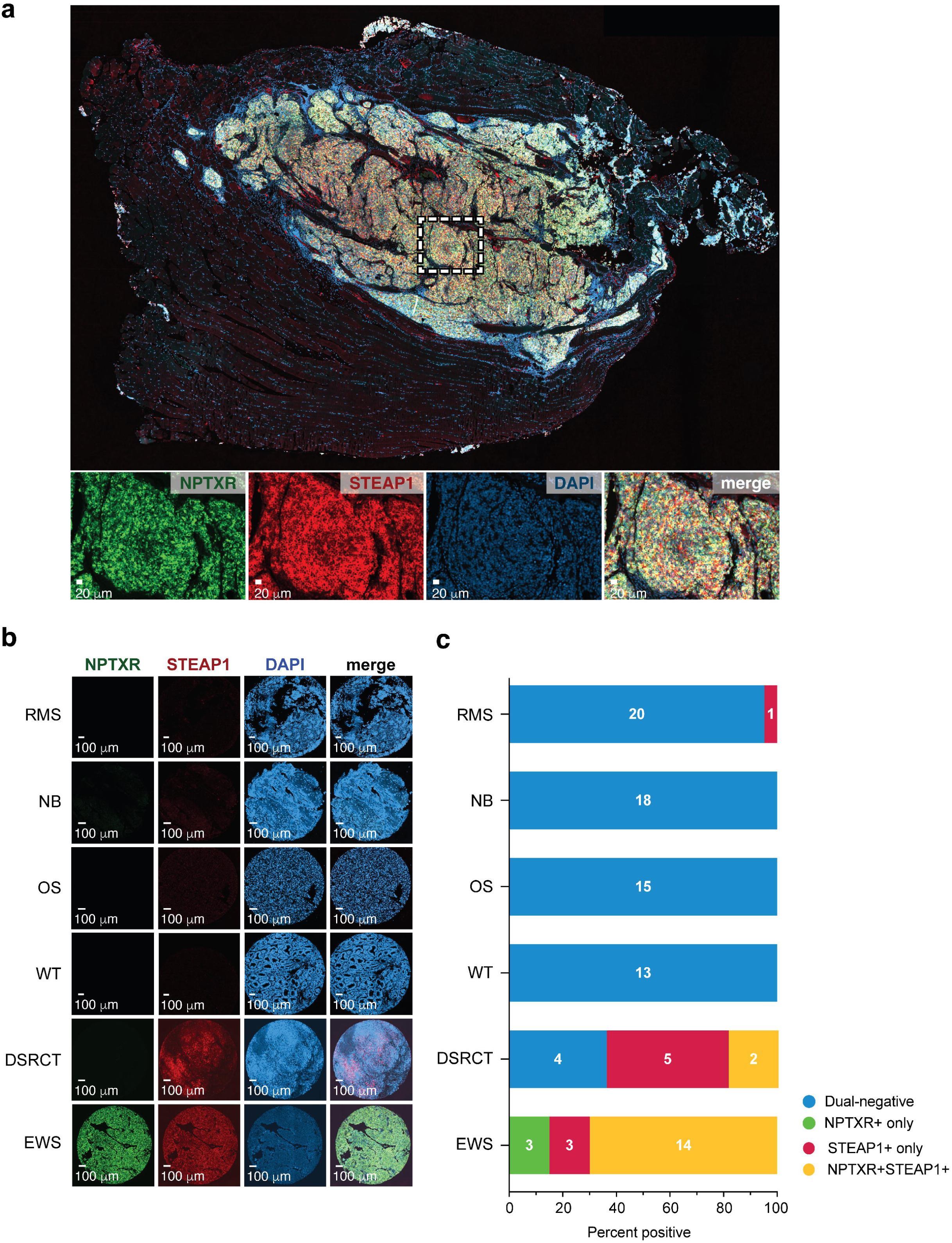
STEAP1 and NPTXR are co-expressed on the surface of Ewing sarcoma cells in O-PDX and patient tumor samples. **a.** Representative dual-immunofluorescence staining of STEAP1 and NPTXR in Ewing sarcoma orthotopic patient-derived xenograft (O-PDX) tumor SJEWS063829_X1. STEAP1 is shown in red, NPTXR in green, and nuclei in blue (DAPI). Images are representative of 4 Ewing sarcoma O-PDX models. Scale bars, 20 μm. **b.** Representative dual-immunofluorescence images of STEAP1 and NPTXR expression in tissue microarrays containing patient tumor samples from Ewing sarcoma (EWS; *n* = 20), desmoplastic small round cell tumor (DSRCT; *n* = 11), Wilms tumor (*n* = 13), osteosarcoma (*n* = 15), neuroblastoma (*n* = 18), and rhabdomyosarcoma (*n* = 21). Scale bars, 100 μm. **c.** Bar plot showing frequency of tumors classified as dual-positive for STEAP1 and NPTXR within each disease group. Additional staining results, sample annotations, and scoring criteria are provided in Supplementary Fig. 6 and Supplementary Table 10.

## DISCUSSION

In this study, we combined a pan-pediatric solid tumor single-cell atlas with a structured AI-based data interpretation agent to address the growing interpretive bottleneck created by massive atlas-scale datasets. By integrating 351 patient and O-PDX datasets across 22 pediatric solid tumor histologies, we recovered 134 robust gene expression programs (GEPs) spanning malignant, immune, stromal, and developmental compartments^4,46^. We then developed an AI agent, based on the Gemini 2.5 LLM, to interpret these GEPs using their gene loadings, functional enrichments, cell-type predictions, malignancy estimates, and deidentified patient metadata. The information provided to the agent, together with its generated annotations and summaries, can be explored and corroborated at pscb.stjude.org. Our approach nominated ectopic expression of neuronal markers in pediatric sarcomas as a cross-disease biological feature with therapeutic relevance, which we further supported through validation of the NPTXR/STEAP1 surface target pair in biologically independent Ewing sarcoma samples.

Our framework addresses a step in single-cell analysis that is distinct from, and complementary to, existing computational methods. NMF-based approaches such as CSI-GEP and single-cell foundation models such as scGPT and scFoundation are designed to extract structure from expression data, yielding cell embeddings, reference-based cell type predictions, or weighted gene modules^1–4^. Converting these outputs into biological understanding has remained a manual, expertise-driven task. Because our agent operates on these outputs rather than on raw expression data, it is agnostic to how GEPs are discovered. In principle, it could be applied downstream of foundation models as readily as downstream of NMF. Importantly, the framework requires no model training, fine-tuning, or dedicated computational infrastructure. All interpretation was performed by querying a general-purpose LLM through a standard API from a laptop computer, and we have provided our prompts. We believe this lowers the barrier for laboratories without specialized AI expertise to extract biological insight from atlas-scale datasets. It also allows the framework to benefit from future improvements in general-purpose LLMs without retraining.

The lineage-mixed programs identified by the agent also suggest a broader set of therapeutic opportunities. By focusing on neurodevelopmental GEPs enriched in Ewing sarcoma, we identified surface proteins that are normally associated with distinct tissue contexts but aberrantly co-expressed on the same malignant cells, with STEAP1/NPTXR illustrating the concept. STEAP1 is a known Ewing sarcoma-associated surface antigen and is of ongoing interest as an immunotherapy target^41,47–49^ with a clinical trial underway (NCT07297979). NPTXR is a neuronal pentraxin receptor with established roles in synaptic organization and it has emerged as an antibody-accessible cancer target in gastric and other cancers^50–52^, but its relevance in Ewing sarcoma has not been reported. STEAP1/NPTXR co-expression in Ewing sarcoma tumors, O-PDX models, and cell lines, together with evidence that normal tissues do not co-express these antigens, nominates this combination as a proof-of-principle candidate for logic-gated therapeutic strategies, including bispecific ADCs or CAR-T. More broadly, these results illustrate a generalizable strategy for target discovery from single-cell atlases by identifying cell states in which two antigens are co-expressed only in malignant cells but are segregated across normal tissue lineages.

Notably, the identification of neuronal and neurodevelopmental programs in pediatric sarcomas should be interpreted in the context of substantial prior literature. Ewing sarcoma has long been recognized as a lineage-ambiguous tumor, historically associated with primitive neuroectodermal features and later linked to mesenchymal or neural crest-like progenitor states^20,21,53–56^. Prior studies have shown that the EWS-FLI1 fusion oncogene can induce Ewing-like transcriptional states, upregulate neural crest-associated genes, and remodel differentiation programs in experimental models^21,55,56^. Similarly, DSRCT is driven by the EWS-WT1 fusion, and EWS-WT1 has been shown to activate neuronal gene expression and partial neural reprogramming^22,57^. Our findings extend this prior work by showing that neuron-like programs can be recovered *de novo* from single-cell transcriptomes, resolved specifically within malignant cells, and detected across a broad spectrum of fusion-driven pediatric sarcomas. The characteristics of these programs in Ewing sarcoma, DSRCT, CIC-rearranged sarcoma, and alveolar rhabdomyosarcoma suggest mosaic activation of developmental regulatory modules rather than faithful recapitulation of a single normal developmental lineage. Notably, these programs were maintained in O-PDX and cell line models, supporting their relevance for functional studies and therapeutic target validation.

One possible explanation for recurrent neurodevelopmental programs across fusion-driven pediatric sarcomas is that oncogenic fusion transcription factors retarget regulatory machinery within permissive developmental chromatin landscapes. In Ewing sarcoma, EWS-FLI1 binds both canonical ETS motifs and GGAA microsatellites, where it can create or remodel enhancer elements and activate disease-specific transcriptional programs^58,59^. EWS-FLI1 also interacts with chromatin-remodeling machinery, including BAF complexes, providing a plausible mechanism for broad enhancer reprogramming^60^. The specific genes activated by these fusions may therefore depend on the cis-regulatory architecture, chromatin accessibility, and cofactor repertoire of the cell of origin or transformation-permissive precursor state. This model could explain why tumors conventionally viewed as mesenchymal can acquire neural, neural crest, glial, or synaptic features without faithfully recapitulating a single normal developmental lineage. More broadly, it suggests that fusion-driven pediatric cancers may be especially prone to non-physiologic combinations of developmental gene expression programs, because their initiating oncogenes directly perturb transcriptional regulation. Mechanistic dissection of these behaviors represents an important direction for future research. For example, perturbation experiments targeting fusion oncogenes, candidate developmental TFs, or surface-antigen programs could determine whether neurodevelopmental GEPs are directly fusion-driven, dependent on specific cofactors, or maintained by additional regulatory circuits

Our approach can be extended in several directions. Expanding the atlas to include additional rare histologies, matched normal pediatric tissues, developmental reference datasets, and spatially resolved profiling would improve disease coverage and strengthen inference about malignant versus normal antigen co-expression. Broader surface-protein validation and functional testing of nominated target pairs will be required to determine which combinations beside *STEAP1*/*NPTXR* should be prioritized for bispecific ADCs, CAR-T cells, or other logic-gated therapeutic strategies. These therapies will ultimately require evaluation in appropriate animal models and clinical trials. Finally, applying a similar strategy to adult tumors could determine whether the lineage-mixed programs described here are specific to pediatric cancers or represent a broader feature of oncogenic transcriptional reprogramming.

In summary, our study shows off-the-shelf LLMs, guided by structured prompting, can serve as a scalable interpretation layer for unsupervised single-cell GEP discovery, recovering known and unexpected developmental programs across pediatric solid tumors. We identified convergent lineage-mixed states across disease boundaries and nominated tumor-selective surface antigen combinations for therapeutic development. More broadly, this work illustrates how AI-guided interpretation of atlas-scale single-cell data can be used to convert large and difficult to analyze biological datasets into experimentally testable and potentially actionable hypotheses.

## METHODS

### Gene program recovery

Integrated single-cell RNA-sequencing (scRNA-seq) data comprising 266 unique pediatric solid tumor samples (351 total datasets including replicates) were analyzed using the scalable unsupervised matrix factorization framework implemented in CSI-GEP^4^. Raw count matrices from all samples were first merged into a unified expression matrix, which was subsequently partitioned into two splits for independent factorization. 5000 highly variable genes were selected prior to factorization^61^.

Briefly, CSI-GEP performs program discovery by using non-negative matrix factorization (NMF) across a wide range of factorization ranks (k = 5 to 500, incremented by 5). For each rank, multiple independent NMF runs (iterations = 100) were conducted using the GPU-accelerated implementation torchnmf, followed by aggregation of results to generate rank-specific consensus matrices. Program stability and reproducibility were evaluated using Jaccard similarity across consensus solutions, computed over multiple Jaccard lengths (ranging from 10 to 100, incremented by 10). The final set of robust gene expression programs was determined by jointly evaluating factorization rank, Jaccard stability metrics, and the number of detected communities. Optimal parameter values were selected through a combination of algorithm-guided criteria and visual inspection of summary curves. In addition, a rescue procedure was applied to retain programs that exhibited moderate stability under relaxed Jaccard similarity thresholds. All downstream analyses were based on this finalized set of programs.

### Prompt engineering and agent design

We developed an automated AI agent based on the Gemini 2.5 large language model to annotate GEPs derived from the PedSTAR atlas. The agent was designed to operate in a stateless manner, so that each GEP was analyzed independently, and annotations for one program did not influence those for others. During development, GPT-5 was additionally used for benchmarking and comparative evaluation. Model interactions were performed primarily through programmatic API calls implemented in custom command-line scripts, with supplementary interactive sessions used for prompt refinement.

For all analyses, we first defined a structured system prompt that provided high-level context and task instructions to the model. This prompt described the nature of the dataset (a pan-pediatric single-cell RNA-seq atlas), defined all input modalities, and specified the analytical objectives and required output format. The system prompt instructed the model to infer the dominant biological function of each GEP, generate a concise descriptive label, provide a short justification grounded in the supplied evidence, and assign a subjective confidence level (High, Moderate, or Low). The expected response structure was explicitly specified to ensure consistent and readable outputs across all programs. The full system prompt is provided in Supplementary File 1.

For each individual GEP, a user prompt was automatically generated using a standardized template. These prompts contained structured information derived from CSI-GEP and downstream analyses. Specifically, we provided the 50 top-ranked genes for each GEP based on NMF loading scores, together with precomputed functional enrichment results. Enrichment analyses were performed using multiple curated gene set collections, including MSigDB Cell Type gene sets, MSigDB Hallmark gene sets, MSigDB positional gene sets (CHR), KEGG pathways, Gene Ontology (Biological Process), transcription factor target gene sets, and immunological gene sets. For each database, significantly enriched terms (false discovery rate < 0.05) were supplied together with enrichment statistics and gene overlaps. In addition, we incorporated cellular context into the user prompts, including SingleR-derived reference-based cell-type prediction scores, NumBat-based malignancy probabilities for individual cells, and metadata describing the disease-of-origin of cells with the highest GEP activity, as defined by GMM thresholding. All inputs were formatted in a structured and standardized manner to ensure consistency across prompts and to minimize ambiguity for the model. A full example of user prompt is provided in Supplementary File 2.

To evaluate how additional contextual information influenced model performance, we tested three independent user prompt configurations. In the first configuration, prompts contained only gene loadings and enrichment results. In the second configuration, SingleR-based cell-type prediction scores were added. In the third configuration, NumBat malignancy probabilities and disease-of-origin metadata were additionally included. Each configuration was applied independently across all GEPs.

Following individual GEP annotation, we performed a second-level meta-annotation analysis. For this step, all AI-generated labels, justifications, confidence scores, and associated metadata across the 134 GEPs were concatenated into a single structured document and provided to the model using a dedicated meta-prompt. The agent was instructed to reason jointly across all programs to identify recurring biological themes, detect co-regulated GEP clusters, and propose experimentally actionable hypotheses. The resulting summary document is provided in Supplementary File 3.

### Cell surface biotinylation assays

Cell surface proteins were selectively enriched by biotin labeling of surface-exposed molecules using the Pierce Cell Surface Protein Isolation Kit (Thermo Scientific, 89881), following the manufacturer’s instructions. The Ewing sarcoma cell line TC-32 was used to confirm plasma membrane molecule expression. Enriched cell surface proteins were then subjected to data-independent acquisition (DIA) mass spectrometry, which was performed by the St. Jude Center for Proteomics and Metabolomics. Briefly, Protein in gel bands were reduced with dithiothreitol (DTT) (Sigma) and alkylated by iodoacetamide (IAA) (Sigma). The gel bands were then washed, dried, and rehydrated with a buffer containing trypsin (Promega). Samples were digested overnight, acidified and the peptides were extracted. The extracts were dried and reconstituted in 5% Formic acid. The peptide samples were loaded on Evotips (Evosep Biosystems) and analyzed on a Bruker timsTOF HT instrument coupled to EvosepOne (Evosep Biosystems) using a 30 samples per day method. Data was acquired using the dia-PASEF mode with an MS1 scan range of 100–1700 m/z. 21 isocratic m/z and ion mobility windows were selected for serial MS2 fragmentation ranging from 475 to 1000 m/z and 0.7 to 1.3 1/K0 respectively. The raw mass spectrometry data from the Bruker HT were searched with DIA-NN (version 2.2.0)^62,63^ using the library-free mode. Protein sequences were downloaded from the Human (Homo sapiens) database (version 2025/09/30). The protein FASTA file was used to build an in silico library with the following settings: digestion enzyme Trypsin/P with maximum 2 missed cleavage; Oxidation on Methionine as variable modification; carbamidomethylated (+57) as fixed modification; maximum number of variable modifications set to 2; peptide length from 7 to 30; precursor charge 1–4; precursor m/z from 400 to 1200; fragment m/z from 200 to 1800. The search parameters of DIA-NN (version 2.2.0) were set as follows: precursor FDR 1%; mass accuracy at MS1 and MS2 both set to 10 ppm; The search results were further filtered with precursor q value <0.01 and protein group q value <0.01 at the library level. For quantification, DIA-NN first obtained precursor quantities by summing the intensities of the top six fragments (ranked by their library intensities) for each precursor. Precursors corresponding to unique proteins were then used for protein-level quantification and intensities of protein groups were obtained using the MaxLFQ algorithm implemented in the iq R package^64^. Enriched cell surface proteins were also quantified by western blot. Input lysate and the biotinylated enriched fraction were loaded at equal protein amounts. Because the non-biotinylated fractions (non-specific control) were processed in parallel from equal cell numbers, they were loaded at equal volumes relative to the biotinylated enriched fraction. Membranes were probed with primary antibodies against STEAP1 (Proteintech, 20199-1-AP, Rabbit Polyclonal antibody, 1:500), NPTXR (Proteintech, 85994-1-RR, 1:500), TOP2A (Cell Signaling Technology, 12286, Rabbit Monoclonal antibody, 1:500), and GAPDH (Cell Signaling Technology, 2118, Rabbit Monoclonal antibody, 1:500), followed by HRP-conjugated anti-rabbit IgG (Cell Signaling Technology, 7074, 1:1000). Signals were detected using Clarity Western ECL Substrate (Bio-Rad, 1705061) and imaged on a ChemiDoc system (Bio-Rad). Band intensities were quantified using Image Lab software (Bio-Rad), and target protein signals were normalized to GAPDH.

### IF staining

Tissues were fixed in 10% neutral buffered formalin, embedded in paraffin, sectioned at 4-μm, mounted on positive charged glass slides (Superfrost Plus; 12-550-15, Thermo Fisher Scientific, Waltham, MA) that were dried at 60°C for 20 minutes, deparaffinized, and stained with hematoxylin and eosin (Richard-Allan Scientific, HE). H&E sections stained and coverslipped using the HistoCore SPECTRA Workstation (Lecia Biosystems). Serial unstained slides were used for multiplex immunofluorescence to co-detect NPTXR (Proteintech, 24807-1-AP, Rabbit Polyclonal antibody, 1:200) and STEAP1 (Proteintech, 20199-1-AP, Rabbit Polyclonal antibody, 1:200).

All staining was performed on the Ventana Discovery Ultra autostainer (Roche, Indianapolis, IN). All tissue sections used for IF labeling underwent deparaffinization followed by heat-induced antigen retrieval using prediluted Cell Conditioning Solution 1 (ULTRA CC1, 950-224, Ventana Medical Systems, Indianapolis, IN). Primary antibodies were serially applied using the U DISCOVERY 5-Plex IF procedure and the following reagents and kits (all from Ventana Medical Systems, Roche): DISCOVERY inhibitor (760-4840), ready to use DISCOVERY OmniMap anti-Rb HRP (760-4311), and visualization of STEAP1 using a DISCOVERY CY5 Kit (760-238) and NPTXR using a DISCOVERY FAM Kit (cat. no. 760-243) and counterstained with DAPI (760-4196). Coverslips were handmounted on slides with Prolong Gold Antifade reagent (Thermo Fisher Scientific, P36931). Sections were digitized with a Zeiss Axiovision Scanner (Zeiss) and imaged using HALO v3.6.4134.137 (Indica Labs, Albuquerque, NM).

## Data and code availability

The Pediatric Solid Tumor Advanced Repository was generated as part of a companion manuscript (McEvoy et al., accepted for publication). That manuscript describes tissue processing, sequencing, and quality control of computational data. We used processed data as Seurat objects, obtained from https://doi.org/10.25452/figshare.plus.31936140. The code and prompt resources used for AI-assisted GEP annotation are available at https://github.com/geeleherlab/PedSTAR. A user-friendly web portal for exploration of the summarized PedSTAR data is available at https://pscb.stjude.org.

## ACKNOWLEDGEMENTS

P.G. is supported by an NIGMS R35 awards [R35GM138293 and R35GM163976] and an R01 grant from NCI [R01CA260060]. A.G.P. has received support from the Rally Foundation, Alex’s Lemonade Stand Foundation, Hyundai Hope on Wheels Foundation, and support through the National Cancer Institute [K08 CA286845]. P.G. and A.G.P. also receive support from ALSAC. The St. Jude Center for Advanced Genome Engineering receives support from NCI [P30CA021765]. Images were acquired at the Cell & Tissue Imaging Center which is supported by SJCRH and NCI [P30CA021765]. The authors thank Sarah K. August for assisting with assembling the text and scientific illustrations. The authors would also like to acknowledge the Comparative Pathology Core Laboratory at St. Jude Children’s Research Hospital.

## Notes

https://doi.org/10.25452/figshare.plus.31936140

https://github.com/geeleherlab/PedSTAR

https://pscb.stjude.org

### Competing Interest Statement

The authors have declared no competing interest.

