## Supplementary Figures for "Lineage-mixed neurodevelopmental programs in pediatric sarcomas nominate tumor-selective surface antigen pairs"

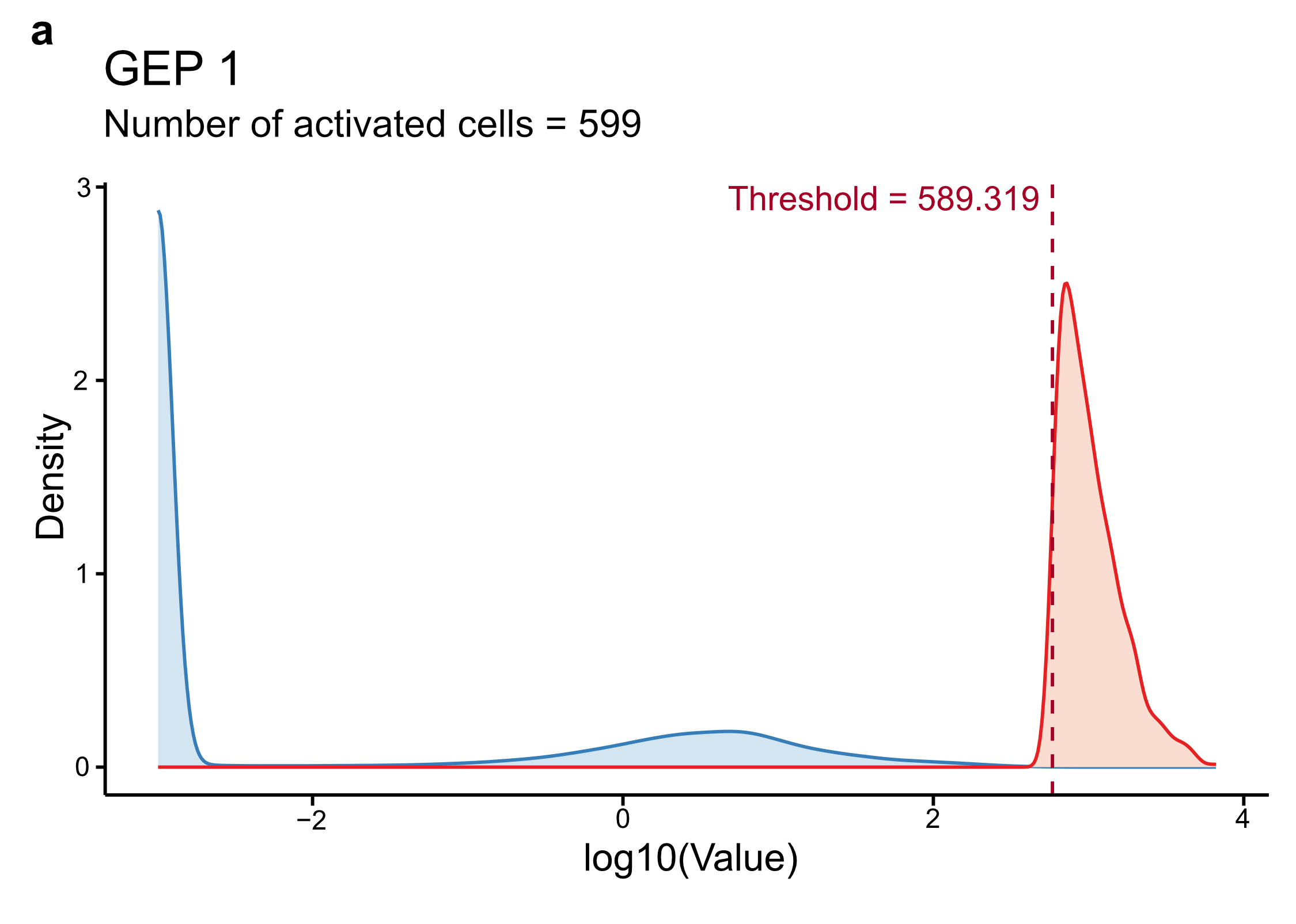

**Supplementary Figure 1**: **Program assignments using GMM.**

Representative density plot of GEP 1 cell activity scores (x-axis) fit with a Gaussian Mixture Model (GMM) to determine whether GEP 1 was active in any given cell. Blue curves indicate the null distribution; the red curve represents the active distribution. The red dashed line marks the calculated activity threshold of 589.319, resulting in 599 GEP1-activated cells. Details of this approach are described in Methods.

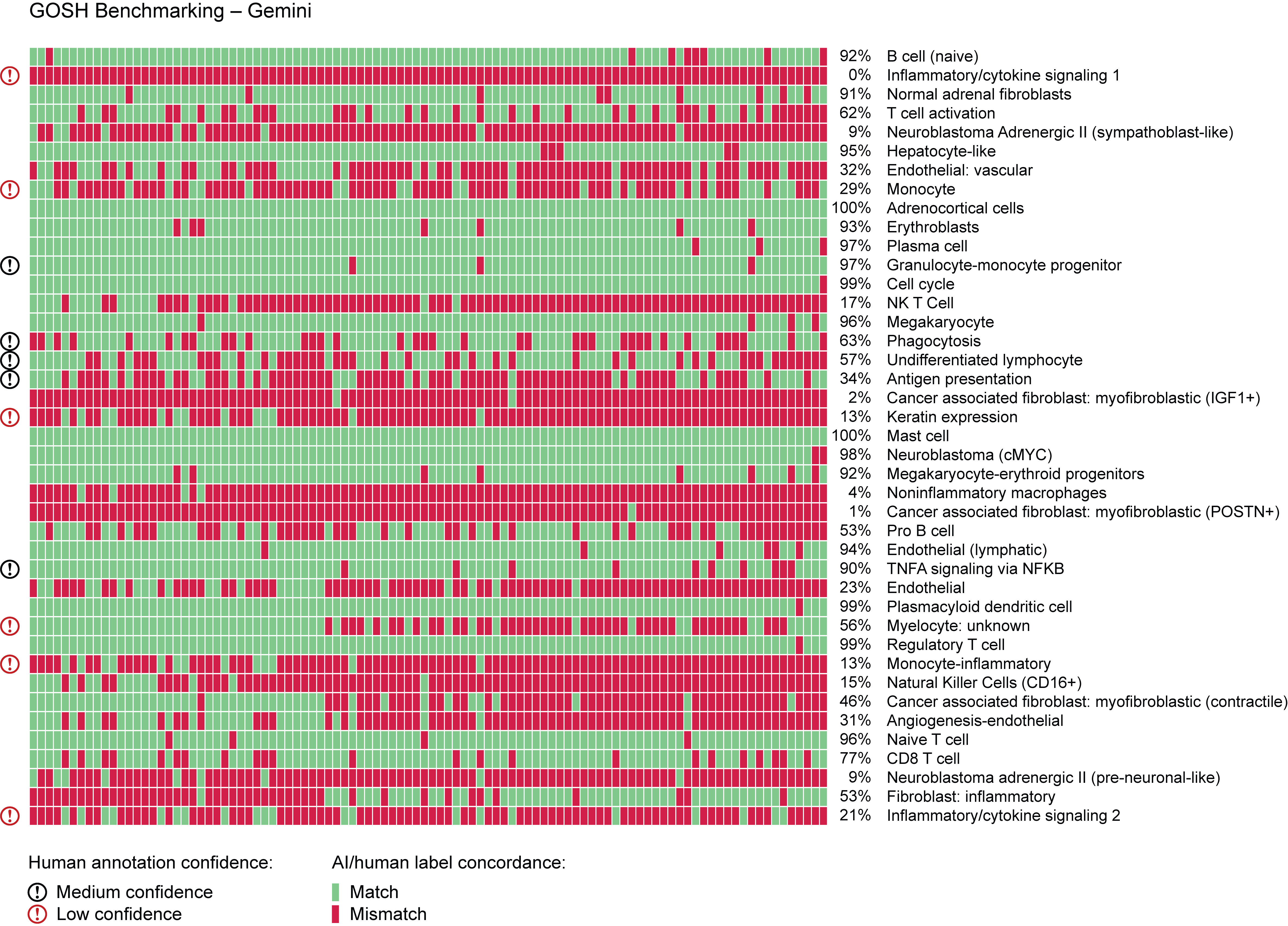

**Supplementary Figure 2: Evaluating the constrained label-matching task of LLMs on a published neuroblastoma dataset previously annotated by human subject matter experts.**

Heatmap showing the Gemini-based AI agent’s matched-to-human-expert (green) and mismatched (red) GEP-label assignments from 100 iterations (columns) for the 41 GEPs (rows) from the GOSH neuroblastoma dataset. Confidence levels from human experts are shown on the left.

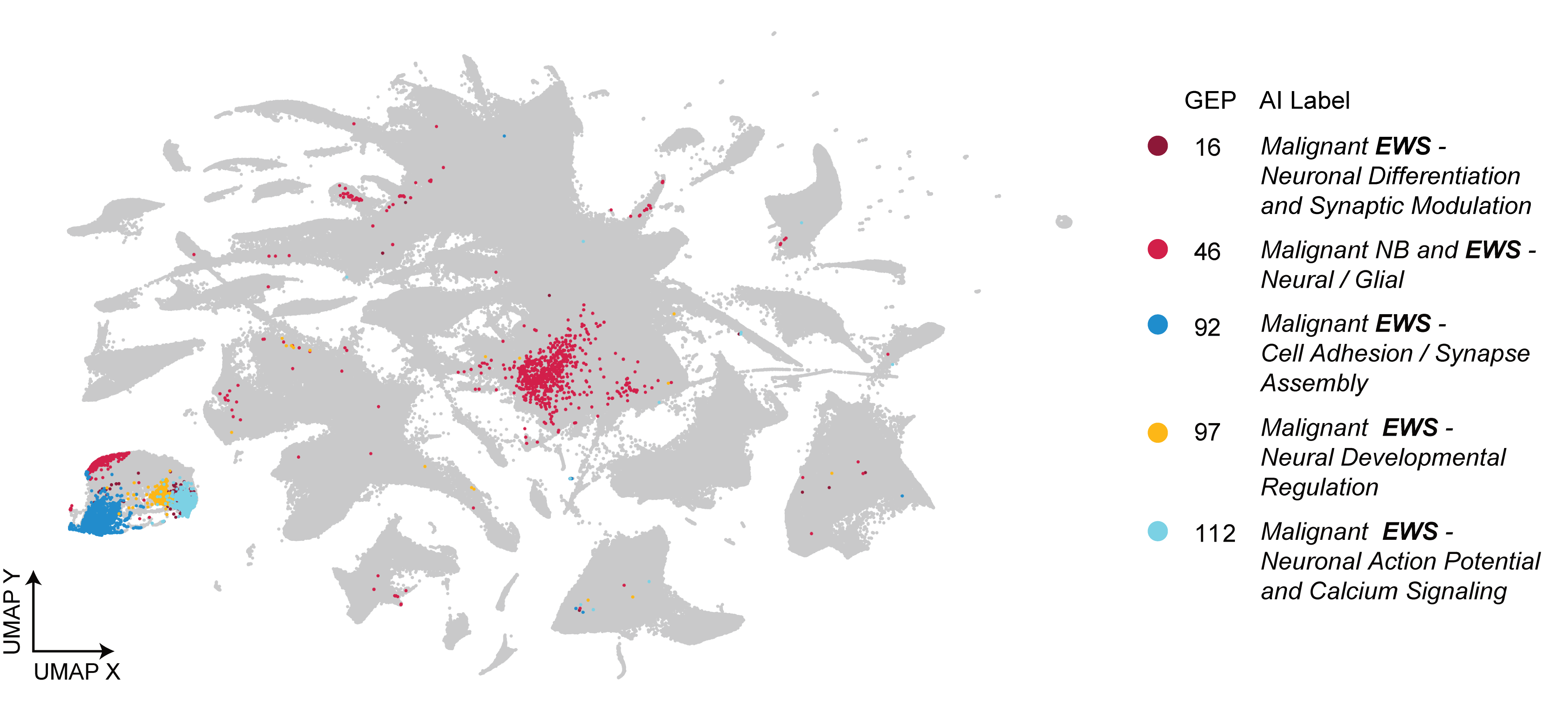

**Supplementary Figure 3: Neurodevelopmental gene expression programs are identified in Ewing sarcoma.**

UMAP projection showing the distribution of Ewing sarcoma (EWS)-specific gene expression programs (GEPs) with neurodevelopmental features across the pediatric cancer atlas. Cells are colored according to activation of individual EWS GEPs, highlighting distinct neurodevelopmental programs associated with neuronal differentiation, synaptic modulation and assembly, neural developmental regulation, and neuronal action potential and calcium signaling.

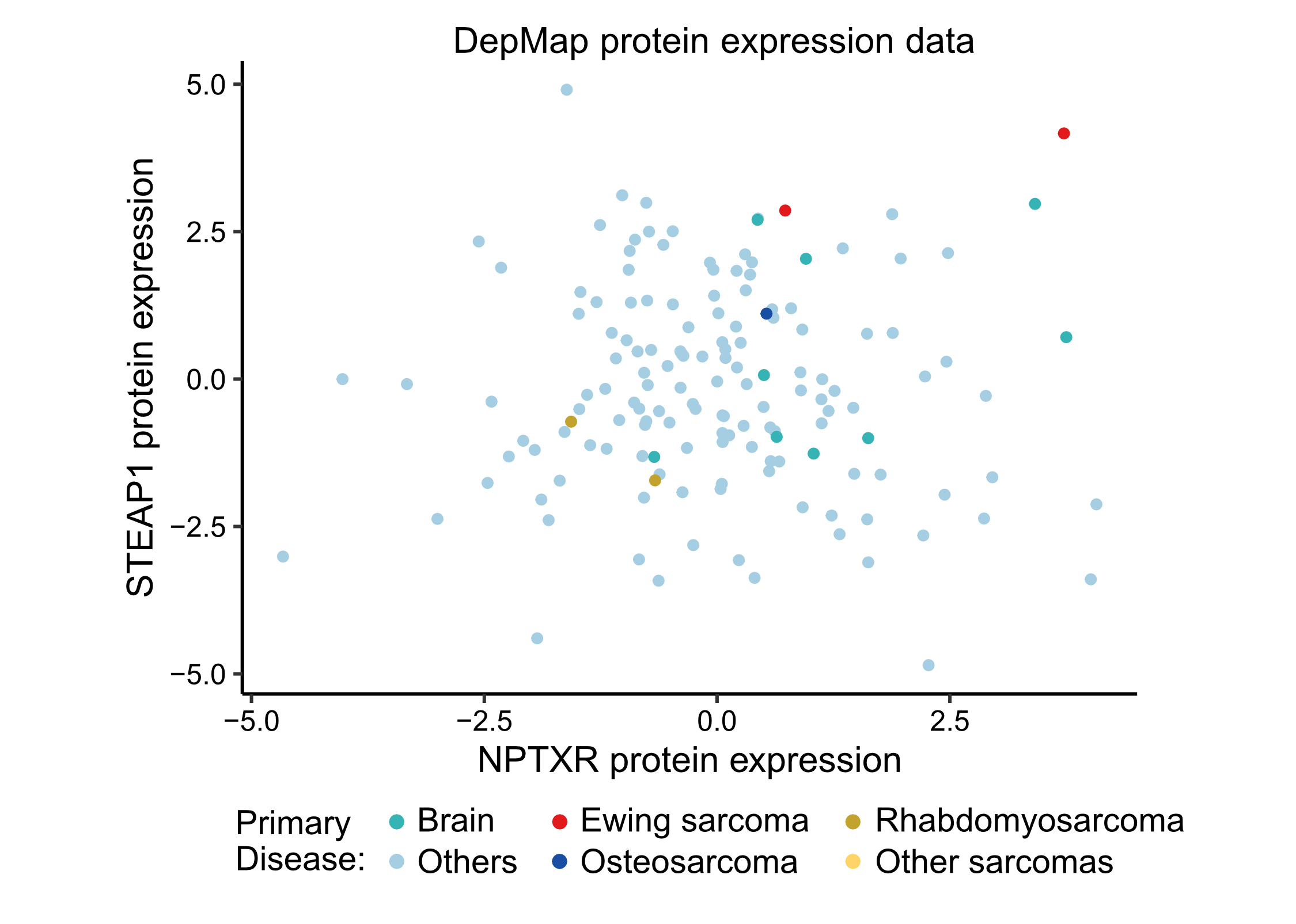

**Supplementary Figure 4: STEAP1 and NPTXR are co-expressed at the protein level in Ewing sarcoma cells.**

DepMap log2 relative protein abundance (from quantitative tandem-mass-tag (TMT) mass spectrometry) showing STEAP1 (y axis) and NPTXR (x axis) expression across cancer cell lines, colored by primary disease. Ewing sarcoma cell lines (red) are among those highly expressing both STEAP1 and NPTXR.
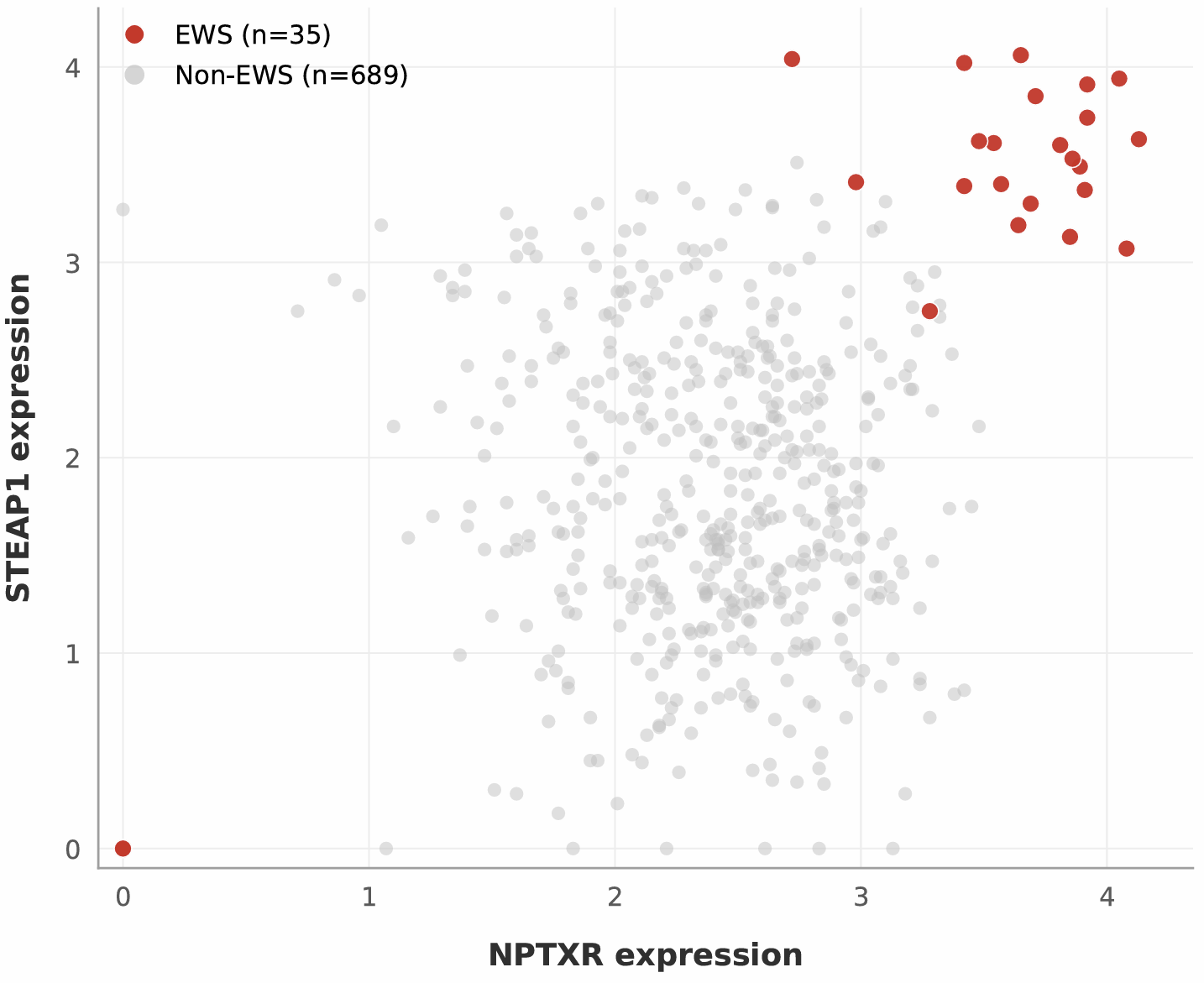

**Supplementary Figure 5: *NPTXR* and *STEAP1* are strongly expressed in Ewing sarcoma patient tumors.**

Bulk RNA-seq log_10_(MoR + 1) expression of NPTXR (x axis) and STEAP1 (y axis) across pediatric solid tumors from the St. Jude PeCan cohort. Ewing sarcoma samples (EWS; *n* = 35, red) are highlighted relative to non-EWS pediatric solid tumors (*n* = 689, gray) and show highly selective overexpression of both genes.

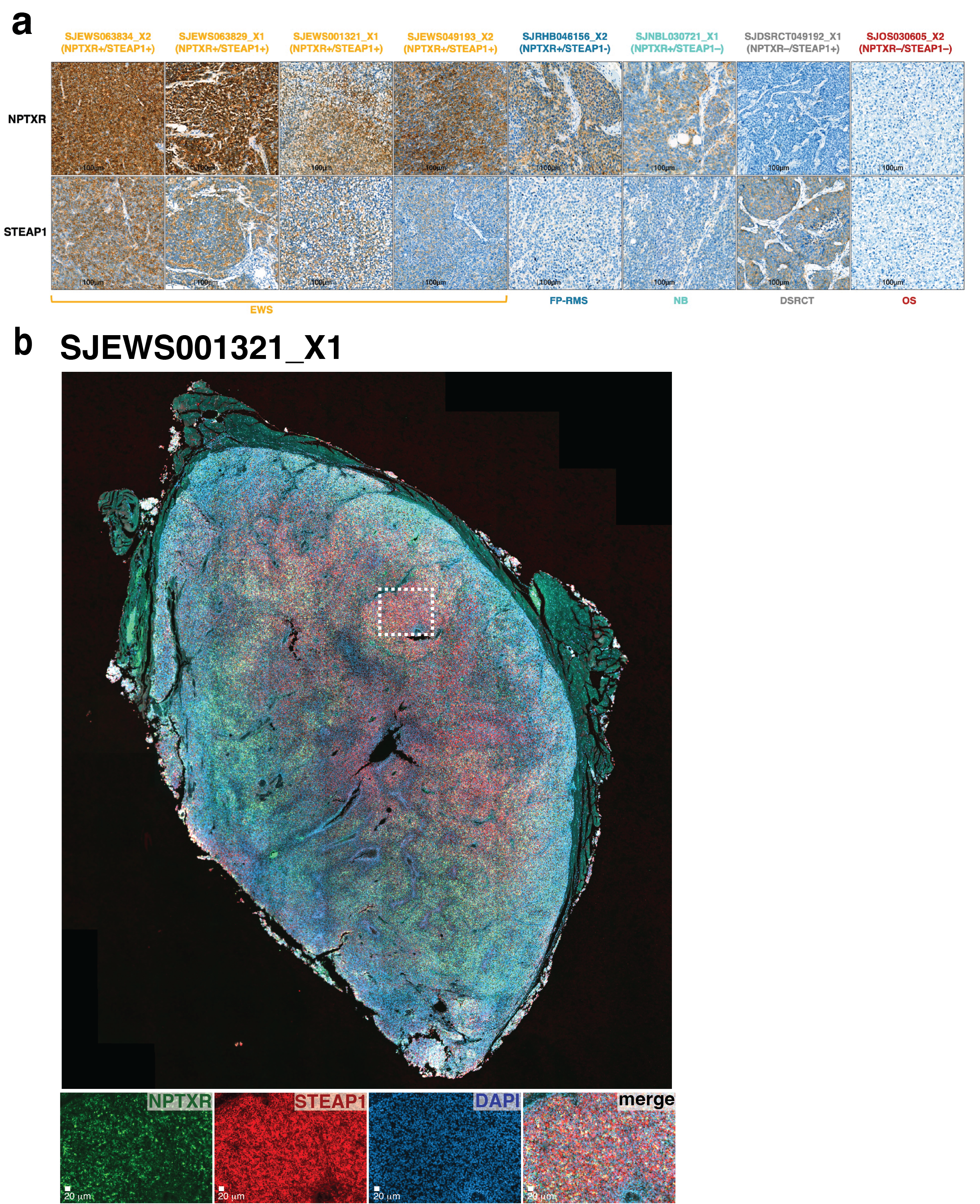

**Supplementary Figure 6: IHC/IF relevant images.**

**a.** Staining of orthotopic xenograft tissue samples from Ewing sarcoma (EWS), fusion-positive rhabdomyosarcoma (FP-RMS), neuroblastoma (NB), desmoplastic small round cell tumor (DSRCT), and osteosarcoma (OS) for NPTXR and STEAP1.

**b.** Dual immunofluorescent staining of a Ewing sarcoma orthotopic xenograft sample, SJEWS001321_X1, for NPTXR and STEAP1. Staining showed dual positivity for both markers with a membranous staining pattern.
