## Supplementary File 1 for "Lineage-mixed neurodevelopmental programs in pediatric sarcomas nominate tumor-selective surface antigen pairs"

**USER PROMPT**

**GEP 1:**

***Top 50 genes with loadings:***

- TM4SF19: 0.0121

- CT69: 0.0075

- TM4SF19-AS1: 0.0068

- LINC01010: 0.0061

- SLC28A3: 0.0053

- KCNA2: 0.005

- CHI3L1: 0.0049

- CHIT1: 0.0046

- CDCP1: 0.0046

- TFEC: 0.0044

- …

- …

**Top enrichment terms:**

From mSigDB Cell Types Gene Set:

- AIZARANI LIVER C6 KUPFFER CELLS 2 | FDR: 1.96e-19 | Genes Found: 15/228

- TRAVAGLINI LUNG TREM2 DENDRITIC CELL | FDR: 6.4e-26 | Genes Found: 23/572

- …

From mSigDB Hallmark Gene Sets:

- HALLMARK COMPLEMENT | FDR: 5.16e-07 | Genes Found: 7/200

- HALLMARK COAGULATION | FDR: 0.0223 | Genes Found: 3/138

- …

From KEGG Pathways:

- KEGG LYSOSOME | FDR: 2.96e-06 | Genes Found: 6/121

- KEGG ANTIGEN PROCESSING AND PRESENTATION | FDR: 0.0224 | Genes Found: 3/88

- …

From CHR Positional Gene Sets:

- chr3q29 | FDR: 0.174 | Genes Found: 3/122

- chr6q23 | FDR: 1 | Genes Found: 2/106

- …

From Transcription Factor Targets:

- NR5A1 TARGET GENES | FDR: 0.0872 | Genes Found: 2/19

- IRF1 01 | FDR: 0.0873 | Genes Found: 4/255

- …

From GO Biological Processes:

- GOBP AMINO SUGAR CATABOLIC PROCESS | FDR: 0.000725 | Genes Found: 3/13

- GOBP CHITIN METABOLIC PROCESS | FDR: 0.0157 | Genes Found: 2/6

- …

From Immunological Gene Sets:

- GSE34156 TLR1 TLR2 LIGAND VS NOD2 AND TLR1 TLR2 LIGAND 24H TREATED MONOCYTE UP | FDR: 4.67e-12 | Genes Found: 11/197

- GSE24634 TREG VS TCONV POST DAY7 IL4 CONVERSION DN | FDR: 4.67e-12 | Genes Found: 11/198

- …

**Number of top cells activating the GEP: 599**

**Mean singleR raw scores across top activated cells:**

- Macrophage: 0.3905

- DC: 0.3598

- Monocyte: 0.3413

- HSC_-G-CSF: 0.2826

- Neutrophils: 0.2726

- Pre-B_cell_CD34-: 0.27

- Platelets: 0.2481

- HSC_CD34+: 0.2455

- NK_cell: 0.2438

- Myelocyte: 0.2374

- T_cells: 0.2322

- GMP: 0.2319

- Pro-Myelocyte: 0.228

- B_cell: 0.228

- BM: 0.2215

- CMP: 0.2144

- Endothelial_cells: 0.2142

- Pro-B_cell_CD34+: 0.208

- Tissue_stem_cells: 0.2028

- Osteoblasts: 0.2026

- Smooth_muscle_cells: 0.2014

- Hepatocytes: 0.1936

- Fibroblasts: 0.1933

- Chondrocytes: 0.1926

- MEP: 0.1896

- Neurons: 0.185

- Epithelial_cells: 0.1823

- iPS_cells: 0.1807

- MSC: 0.1737

- BM & Prog.: 0.1714

- Erythroblast: 0.1682

- Astrocyte: 0.1622

- Keratinocytes: 0.1602

- Gametocytes: 0.151

- Embryonic_stem_cells: 0.134

- Neuroepithelial_cell: 0.114

**Disease origin for top activated cells:**

- ERMS: 0.8331

- NB: 0.1569

- RB: 0.0067

- EWS: 0.0017

- GCT: 0.0017

**Mean NumBat probability of malignancy across top activated cells:**

- SNV: 0.037 (sd: 0.143)

- CNV: 0.37 (sd: 0.482)

- Joint: 0.217 (sd: 0.409)
