## Supplementary File 3 for "Lineage-mixed neurodevelopmental programs in pediatric sarcomas nominate tumor-selective surface antigen pairs"

**SYSTEM PROMPT**

You are a molecular biology expert leading a translational research laboratory that integrates computational biology and experimental molecular biology. You are analyzing **134 gene expression programs (GEPs)** obtained via non-negative matrix factorization (**NMF**) of single-cell RNA-seq data from a pan-cancer pediatric cohort.

You are provided with:

- **GEP Annotation Table**: This table describes each GEP, including functional labels, biological justifications, and confidence levels.

- **GEP Co-expression Matrix**: A matrix of pairwise Spearman's RHO coefficients and p-values between GEPs.

Task:

**Based on these data, propose comprehensive and well-justified experimental validation plans**, addressing the following:

- Scientific **Rationale and Novelty**

  - For each proposed experiment, explain the novelty of the observation you intend to validate.

  - Justify why it is biologically relevant and interesting, particularly in the context of pediatric cancers.

- Use the **GEP co-expression** matrix to underline the interest of the observation to validate.

- Compartment-Specific Strategies

  - Propose validation strategies tailored to:

    - **Malignant (Tumor)** Compartment, for example functional assays, perturbation studies.

   - **Immune** Compartment, for example immune profiling, cytokine analyses, cell-cell interactions.

   - **Stromal** Compartment, for example fibroblast characterization, extracellular matrix analyses.

For each compartment:

    - Explain the **biological hypothesis**, the **experimental approach**, the expected **outcomes** and their implications.

  - Prioritize the use of **cutting-edge technologies**

  - Clearly describe which technology to use for each experiment and why.

Update:

Provide experimental plans specific to each disease identified in the dataset.

**AI-Agent (Gemini 2.5-based) OUTPUT:**

**Comprehensive Experimental Validation Plans for Pediatric Cancer Gene Expression Programs**

**I. Executive Summary**

This report delineates comprehensive experimental validation plans for 134 gene expression programs (GEPs) identified through non-negative matrix factorization (NMF) of single-cell RNA-seq data from a pan-cancer pediatric cohort. These GEPs represent distinct biological processes within malignant, immune, and stromal compartments of pediatric tumors. The proposed strategies are designed to leverage cutting-edge technologies, including spatial transcriptomics and patient-derived xenografts (PDX), to confirm novel biological observations, elucidate functional interactions between GEPs, and identify potential diagnostic, prognostic, and therapeutic targets. A particular focus is placed on Neuroblastoma and Osteosarcoma, given their unique biological characteristics and the prevalence of specific GEPs within these tumor types. The ultimate aim is to translate these computational discoveries into actionable insights for improved pediatric cancer treatment.

**II. Introduction: Unveiling Pediatric Cancer Biology through GEPs**

Single-cell RNA sequencing (scRNA-seq) has fundamentally transformed the landscape of cancer research by providing an unprecedented resolution of gene expression profiles at the individual cell level. This technological advancement has been instrumental in dissecting the complex cellular heterogeneity inherent in tumors, moving beyond bulk tissue analyses that often obscure critical cell-specific signals. Non-negative matrix factorization (NMF) stands as a powerful computational method for deconvoluting these intricate scRNA-seq datasets. By applying NMF, complex gene expression patterns are resolved into a smaller set of interpretable GEPs, each representing a co-regulated network of genes that defines specific cellular states or underlying biological processes within the tumor microenvironment.

The significance of identifying these GEPs extends to providing a more granular yet comprehensive understanding of the diverse cellular landscapes within pediatric cancers. Each GEP offers a snapshot of a particular biological program, whether it pertains to the intrinsic characteristics of malignant cells, the functional states of infiltrating immune cells, or the supportive and remodeling activities of stromal components. By dissecting these distinct programs, researchers can begin to unravel the intricate interplay between different cell types that collectively drives tumor initiation, progression, metastasis, and, critically, resistance to therapy. This detailed, cell-centric view is indispensable for moving towards precision medicine in pediatric oncology, where treatments can be tailored to the specific molecular vulnerabilities and microenvironmental dependencies of a child's tumor.

While computational identification of GEPs provides robust hypotheses regarding their biological roles, experimental validation is an indispensable step to confirm their relevance, functional impact, and translational potential. This is particularly crucial in the context of pediatric cancers, which often present with unique developmental contexts, distinct molecular drivers, and a pressing need for novel therapeutic strategies due to limited treatment options and the long-term side effects of conventional therapies. Furthermore, the scarcity of patient material in pediatric oncology necessitates highly targeted and efficient research strategies. Validating these computationally derived GEPs through rigorous experimental approaches will transition observations from correlative associations to mechanistic understanding, thereby paving the way for the development of novel diagnostic biomarkers, more accurate prognostic indicators, and ultimately, more effective and less toxic therapeutic interventions for children with cancer.

**III. Scientific Rationale and Novelty of GEP Validation**

**Overview of Key GEPs for Validation**

The 134 identified GEPs, all designated with "High" confidence in the GEP Annotation Table ^1^, represent robust computational derivations that warrant thorough experimental scrutiny. These programs span a diverse array of biological categories, offering a rich resource for understanding pediatric cancer biology. A significant proportion of these GEPs fall under the "Neurodevelopmental" category, with many exhibiting high specificity to Neuroblastoma (NB), such as Program 7 ("Malignant NB - Neuronal Proliferation and Cholinergic Signaling"), Program 15 ("Malignant NB - Neuronal Synaptic and Calcium Signaling"), Program 33 ("Malignant NB - Core Oncogenic/Proliferative - chr2p24 amplification"), Program 40 ("Malignant NB - Neuroendocrine and Neuronal Differentiation"), Program 54 ("Malignant NB - Neuronal Ion Homeostasis and Signaling"), Program 58 ("Malignant NB - Mitochondrial Respiration and Axonal Transport"), Program 107 ("Malignant NB - Synaptic Vesicle Dynamics and Neuronal Excitability"), Program 109 ("Malignant NB - Synaptic Plasticity and Neuropeptide Signaling"), Program 110 ("Malignant NB - Axonogenesis and Cholinergic Receptor"), and Program 111 ("Malignant NB - Neuronal Axon Guidance").^1^ Ewing Sarcoma (EWS) also exhibits specific neurodevelopmental programs, including Program 16 ("Malignant EWS - Neuronal Differentiation and Synaptic Modulation"), Program 92 ("Malignant EWS - Cell Adhesion/Synapse Assembly"), Program 97 ("Malignant EWS - Neural Developmental Regulation"), and Program 112 ("Malignant EWS - Neuronal Action Potential and Calcium Signaling").^1^

Beyond malignant programs, the dataset includes numerous "Immune and Inflammatory Response" GEPs, characterizing various immune cell types and their functional states within the tumor microenvironment. Examples include Program 1 ("Non-malignant Macrophage / Myeloid Activation"), Program 17 ("Non-malignant Macrophage / Osteoclast Differentiation"), Program 22 ("Non-malignant Humoral Immune Response"), Program 57 ("Non-malignant T Cell Receptor Signaling"), Program 67 ("Non-malignant Neutrophil Activation"), Program 68 ("Non-malignant Dendritic Cell-Like / Antigen Presentation"), Program 69 ("Non-malignant Macrophage / Monocyte-Like"), Program 77 ("Non-malignant NK Cell Cytotoxicity and Chemotaxis"), Program 87 ("Non-malignant Cytotoxic T Cell and NK Cell Effector"), Program 108 ("Non-malignant T Cell Activation"), Program 119 ("Non-malignant Interferon-Gamma Response"), Program 121 ("Non-malignant B Cell and Plasma Cell Immune Response"), Program 129 ("Non-malignant Interferon Response"), and Program 134 ("Non-malignant Myeloid Cell / Neutrophil Innate Immunity").^1^ A key "Stromal and Tissue Remodeling" program, Program 2 ("Malignant OS - Bone Morphogenesis and Extracellular Matrix"), is specifically associated with Osteosarcoma.^1^ The primary focus of experimental validation will be on malignant GEPs that define tumor-specific characteristics and non-malignant GEPs that characterize the tumor microenvironment,Yes, especially those with high disease specificity or strong co-expression patterns.

**Biological Relevance in Pediatric Cancers**

Validating these GEPs is of paramount importance for advancing the understanding of the unique biology underpinning pediatric cancers. For instance, the striking prevalence of "Neurodevelopmental" GEPs in Neuroblastoma underscores the disease's neural crest origin and its profound reliance on aberrantly activated developmental pathways for sustained proliferation and survival. Confirming the functional roles of specific genes within these GEPs, such as *MYCN* (a known oncogene) in Program 7 and 33, or *ALK* in Program 15 and 33, can illuminate critical oncogenic dependencies that may serve as therapeutic vulnerabilities.^1^

Similarly, deciphering the precise roles of specific immune programs, such as Program 87 ("Non-malignant Cytotoxic T Cell and NK Cell Effector") in Neuroblastoma, or the stromal programs like Program 2 ("Malignant OS - Bone Morphogenesis and Extracellular Matrix") in Osteosarcoma, is crucial. These programs represent the intricate interactions within the tumor microenvironment that dictate crucial aspects of tumor biology, including immune evasion, metastatic dissemination, and resistance to conventional therapies.^1^ Experimental validation will move beyond mere correlation to establish causal relationships, providing a mechanistic foundation for developing novel strategies to manipulate the tumor or its microenvironment for therapeutic benefit.

**Leveraging GEP Co-expression for Novelty**

The GEP co-expression matrix provides a powerful lens through which to identify novel biological observations that warrant in-depth experimental validation. High Spearman's RHO coefficients, particularly those with associated p-values of 0, indicate statistically significant and strong co-regulation or synergistic functions between GEPs.^1^ These highly correlated pairs represent compelling new avenues for investigation.

One notable observation is the coordinated nature of malignant Neuroblastoma identity. The co-expression matrix reveals several top co-expressed GEP pairs involving malignant Neuroblastoma programs. For example, Program 15 ("Malignant NB - Neuronal Synaptic and Calcium Signaling") and Program 110 ("Malignant NB - Axonogenesis and Cholinergic Receptor") demonstrate a high positive correlation (Rho = 0.862).^1^ Similarly, Program 33 ("Malignant NB - Core Oncogenic/Proliferative - chr2p24 amplification") and Program 111 ("Malignant NB - Neuronal Axon Guidance") exhibit a strong correlation (Rho = 0.855).^1^ Program 54 ("Malignant NB - Neuronal Ion Homeostasis and Signaling") and Program 109 ("Malignant NB - Synaptic Plasticity and Neuropeptide Signaling") are also highly correlated (Rho = 0.862).^1^ A review of the GEP annotations confirms that all these programs are explicitly designated as "Malignant NB" and consistently describe themes of neuronal development, signaling, axon guidance, and proliferation.^1^ Program 33, along with Program 7 (which is highly correlated with GEP 117 at 0.966, though GEP 117's description is not provided), specifically highlights the presence of

*MYCN* and *chr2p24 amplification*, which are well-established oncogenic drivers in Neuroblastoma.^1^ The exceptionally high positive correlations observed among these malignant Neuroblastoma-specific neuronal GEPs strongly suggest that they do not function as isolated biological processes. Instead, they appear to represent a tightly integrated and co-regulated network that collectively defines the core malignant neuronal identity and developmental state of Neuroblastoma cells. This implies a coordinated oncogenic program that orchestrates multiple aspects of tumor biology, rather than a collection of independent pathways. If these GEPs indeed function as a unified malignant program, perturbing one component might have cascading effects across the entire network. This understanding opens up promising avenues for identifying master regulators—such as key transcription factors or central signaling hubs like ALK or MYCN, which are identified as top genes in some of these GEPs ^1^—that exert control over multiple facets of Neuroblastoma malignancy. Targeting such master regulators could offer more effective and durable therapeutic strategies by disrupting a broader malignant network, potentially overcoming resistance mechanisms that arise from targeting single pathways. This also indicates that the specific neuronal differentiation state of Neuroblastoma cells is intrinsically linked to their proliferative and aberrant signaling characteristics.

Another significant finding pertains to the convergent neuronal phenotypes observed in sarcomas. The co-expression matrix highlights a strong correlation (Rho = 0.890) between Program 16 ("Malignant EWS - Neuronal Differentiation and Synaptic Modulation") and Program 112 ("Malignant EWS - Neuronal Action Potential and Calcium Signaling"), both annotated as Ewing Sarcoma-specific malignant programs.^1^ Despite Ewing Sarcoma being a mesenchymal tumor, GEPs 16 and 112 explicitly describe prominent neuronal features, including differentiation, synaptic modulation, action potentials, and calcium signaling.^1^ Furthermore, the GEP annotations reveal similar neuronal characteristics in other sarcoma types: Program 14 ("Malignant HGS - Neuronal Signaling and Development"), Program 62 ("Malignant DSRCT - Neuronal and Astrocytic Development"), Program 82 ("Malignant DSRCT- Neuronal Adhesion / Synapse Organization"), and Program 118 ("Malignant ARMS - Myogenesis / Neural Development").^1^ The consistent emergence of distinct, yet functionally related, neuronal GEPs across various pediatric sarcomas (Ewing Sarcoma, High-Grade Sarcoma, Desmoplastic Small Round Cell Tumor, Alveolar Rhabdomyosarcoma) points to a significant phenomenon of convergent oncogenic differentiation or developmental plasticity. This is a non-obvious finding because these tumors originate from diverse mesenchymal lineages, yet they appear to aberrantly activate common neuronal developmental programs. This challenges the conventional understanding of these tumors as purely mesenchymal entities. This shared "neuronal-like" vulnerability across different sarcoma types could lead to the identification of pan-sarcoma therapeutic targets. For example, if calcium signaling (highlighted in Program 112) is a common dependency in these neuronal-like sarcomas, then calcium channel modulators or related signaling inhibitors could potentially be broadly effective across these distinct tumor types. Moreover, understanding the molecular mechanisms driving this aberrant neuronal differentiation could reveal novel therapeutic strategies that aim to force terminal differentiation or reverse the malignant phenotype by disrupting these developmental programs.

A critical question arises regarding the interpretation of "malignant" labels in certain immune/stromal GEPs. For instance, Program 9 is labeled "Malignant NB - B Cell-Like" and Program 26 is labeled "Malignant NB - Platelet / Megakaryocyte-Like" in the GEP Annotation Table.^1^ However, a closer examination of their "NumBat malignancy probabilities" reveals that they are not "very high" (Program 9: Joint 0.648; Program 26: Joint 0.464).^1^ Crucially, their descriptions explicitly state that Program 9 is "likely driven by B cell infiltration" and Program 26 "strongly reflects platelet activation and megakaryocyte biology".^1^ This contrasts with other immune GEPs like Program 22 and 121 (both B/Plasma cell related), which are explicitly interpreted as

*non-malignant* based on similar NumBat scores.^1^ There is a discernible nuance or potential contradiction in the labeling and interpretation of malignancy for these GEPs. The "Malignant" label for Programs 9 and 26 might refer to their strong association with malignant tumors, or perhaps a very subtle, partial malignant transformation within a small subset of cells. However, the descriptive text and NumBat scores lean towards them representing non-malignant infiltrating immune/hematopoietic cells. This highlights a critical unresolved question within the dataset that necessitates rigorous experimental validation to definitively classify these GEPs. It is paramount to distinguish between tumor-intrinsic programs that mimic immune/stromal cell states (e.g., through aberrant differentiation or lineage plasticity of tumor cells) and genuine infiltration by non-malignant immune/stromal cells. If tumor cells themselves are expressing these GEPs, they represent direct therapeutic targets for differentiation therapy or lineage-specific cytotoxic agents. Conversely, if they are primarily infiltrating immune cells, the therapeutic strategy would shift towards modulating the tumor microenvironment (e.g., enhancing anti-tumor immunity or depleting pro-tumorigenic immune cells). This distinction has profound and direct implications for the design of clinically translatable therapeutic strategies.

Furthermore, the data suggests metabolic reprogramming in pediatric cancers that extends beyond the classic Warburg effect. Program 58 ("Malignant NB - Mitochondrial Respiration and Axonal Transport") explicitly lists "KEGG OXIDATIVE PHOSPHORYLATION" and includes mitochondrial genes such as MT-ATP8, MT-ND3, and MT-ND2.^1^ Similarly, Program 88 ("Malignant RB - Retinal Neurodevelopment and Oxidative Phosphorylation") highlights "HALLMARK OXIDATIVE PHOSPHORYLATION" and features mitochondrial genes like SLC25A4, UQCRHL, and COX8A.^1^ Both GEPs are described as high-confidence malignant programs, specific to Neuroblastoma and Retinoblastoma, respectively, both of which are pediatric cancers.^1^ The prominent and recurrent enrichment of oxidative phosphorylation (OXPHOS) pathways and mitochondrial genes in these distinct malignant pediatric neuronal tumors is a significant finding. While the Warburg effect (aerobic glycolysis) is a classic metabolic hallmark of rapidly proliferating cancer cells, this data indicates that some pediatric cancers, particularly those with a strong neuronal phenotype, may rely substantially on mitochondrial respiration for energy production and biosynthesis. This observation challenges a simplistic, universal view of cancer metabolism, indicating metabolic heterogeneity even within pediatric tumors. This reliance on OXPHOS opens a novel therapeutic window for these specific pediatric cancers. Targeting OXPHOS, for instance, with mitochondrial inhibitors or by disrupting key components of the electron transport chain, could represent a potent and specific therapeutic strategy. This warrants further in-depth investigation into the metabolic vulnerabilities of these particular tumor types, potentially leading to the development of new drugs or the repurposing of existing metabolic modulators for pediatric oncology.

Finally, the analysis points to significant developmental plasticity and mixed-lineage states in pediatric sarcomas. Program 118 ("Malignant ARMS - Myogenesis / Neural Development") is specific to Alveolar Rhabdomyosarcoma (ARMS) and explicitly shows both myogenic and neural developmental features, including the muscle-related gene TNNT2 and neuronal markers like NOS1 and DCX.^1^ Program 120 ("Malignant NB - Neuroendocrine / Cartilage Development"), while broadly distributed, is prevalent in Neuroblastoma and displays characteristics of both cartilage development (e.g., collagens, ACAN) and neuroendocrine features (e.g., PROK1, VGF, CRH, VIP).^1^ Furthermore, Program 74 ("Malignant URCS - Mesenchymal and Neural Development") integrates kidney epithelial and neural developmental characteristics.^1^ All these GEPs are identified as malignant programs arising from diverse pediatric tumor types (ARMS, NB, URCS).^1^ The presence of these "mixed-lineage" or "developmental plasticity" GEPs represents a profound biological finding into the fundamental nature of pediatric sarcomas. It suggests that these tumors, often originating from primitive or progenitor cells, retain significant developmental plasticity. This allows them to aberrantly express gene programs characteristic of multiple embryonic lineages, indicating complex and often incomplete or misdirected developmental trajectories. This phenomenon goes beyond simple dedifferentiation and points to a more intricate biological underpinning of tumor heterogeneity. This inherent developmental plasticity could be a major contributing factor to tumor heterogeneity, therapeutic resistance, and metastatic potential in pediatric cancers. Understanding the precise regulatory networks that enable this plasticity (e.g., Hedgehog signaling, mentioned in Program 120, which is a key developmental pathway) could identify novel therapeutic targets. These targets might aim to force terminal differentiation of tumor cells or block these aberrant developmental pathways, thereby sensitizing tumor cells to conventional therapies or preventing disease relapse. This also highlights the crucial need for multi-lineage targeting strategies to effectively address the diverse cellular states within these complex tumors.

**IV. Compartment-Specific Experimental Validation Strategies**

This section details experimental plans tailored to the malignant, immune, and stromal compartments, emphasizing the integration of cutting-edge technologies to validate specific biological hypotheses.

**A. Malignant (Tumor) Compartment**

**Scientific Rationale and Novelty**

Malignant GEPs are hypothesized to reflect distinct tumor cell states, encompassing critical processes such as proliferation, differentiation, stemness, and drug resistance. These programs are often driven by specific oncogenic pathways or aberrant developmental programs, which are fundamental for tumor maintenance and progression in pediatric cancers. The novelty lies in validating the functional impact of these computationally identified GEPs on tumor cell behavior and identifying their underlying molecular drivers within the specific context of pediatric malignancies.

**Experimental Approach**

To directly assess the phenotypic consequences of GEP activation or inhibition on tumor cell behavior, a multi-faceted experimental approach is proposed:

1. **Functional Assays in 2D/3D Culture Systems**:
   - **Design**: Representative malignant GEPs will be selected for validation. For Neuroblastoma, this includes programs such as Program 7, 15, 33, 40, 54, 58, 107, 109, 110, and 111, which are characterized by neuronal development, proliferation, and signaling.^1^ For Ewing Sarcoma, Program 16, 112, 92, and 97, which highlight neuronal differentiation and cell adhesion, will be prioritized.^1^ For Osteosarcoma, Program 2, focusing on bone morphogenesis, is a key candidate.^1^ Pediatric cancer cell lines (e.g., Neuroblastoma: SK-N-BE(2), SH-SY5Y; Ewing Sarcoma: A673, TC-71; Osteosarcoma: U2OS, Saos-2) that endogenously express these GEPs at varying levels will be utilized. Following modulation of GEP activity (e.g., via gene perturbation or small molecule inhibitors), assays will be performed to quantify:
     - **Proliferation**: Measured by MTT/alamarBlue assays, EdU incorporation, and direct cell counting to determine cell growth rates.
     - **Migration/Invasion**: Assessed using Transwell assays and scratch assays to evaluate the metastatic potential of tumor cells *in vitro*.
     - **Differentiation**: Induction of specific lineage markers will be examined. For Neuroblastoma and Ewing Sarcoma GEPs, this includes neuronal markers such as neurite outgrowth and expression of SYN3, MAP2 (from GEPs 54, 107, 109).^1^ For Osteosarcoma GEP 2, osteogenic markers like alkaline phosphatase activity and calcium deposition will be assessed.
     - **Spheroid/Organoid Formation**: The use of 3D culture models will enable the assessment of self-renewal capacity, stemness, and the ability to form tissue-like structures, which is particularly relevant for neurodevelopmental GEPs and their differentiation potential.
   - **Technology Integration**: High-content imaging platforms will be employed for automated, quantitative analysis of phenotypic changes. Time-lapse microscopy will provide dynamic insights into cellular processes such as migration and neurite extension.
2. **Perturbation Studies (CRISPR/Cas9, shRNA/siRNA, Overexpression)**:
   - **Rationale**: To establish causal links between specific GEPs (or their key driver genes) and the observed malignant phenotypes.
   - **Design**: Based on the top genes identified in the GEP descriptions ^1^, key genes within selected malignant GEPs will be targeted for perturbation. For example,

*MYCN* and *ALK* in Neuroblastoma Programs 7 and 33; *NTRK3* in Desmoplastic Small Round Cell Tumor Program 82; *SOX6*, *PAX7*, *PAX3*, and *NKX2-2* in Ewing Sarcoma Program 16; *TNNT2* and *NOS1* in Alveolar Rhabdomyosarcoma Program 118; and *ACAN* in Osteosarcoma Program 2.^1^ CRISPR/Cas9 will be used for precise gene editing (knockout or activation/inhibition), while shRNA/siRNA will facilitate gene knockdown. Overexpression constructs will be employed to investigate gain-of-function effects.

3. **Technology Integration for Malignant Compartment**:
   - **Patient-Derived Xenografts (PDX)**: PDX models are critical for *in vivo* validation of GEPs and their functional roles in tumor growth, metastasis, and response to therapeutic interventions. These models faithfully recapitulate the tumor's genetic and histological heterogeneity, as well as aspects of the tumor microenvironment, making them superior to conventional cell line xenografts for translational studies. PDX will be used to confirm the impact of GEP perturbations on tumor progression and to evaluate the efficacy of GEP-targeted therapies *in vivo*.
   - **Spatial Transcriptomics**: This cutting-edge technology will be instrumental in mapping the precise expression patterns of malignant GEPs within the tumor architecture. By correlating GEP activity with histological features, cellular morphology, and the spatial distribution of neighboring cells, it will be possible to understand how GEPs contribute to distinct tumor niches and how they interact with other cellular compartments *in situ*. This will provide crucial context that is lost in dissociated single-cell data.

**Expected Outcomes and Their Implications**

These experiments are expected to yield several key outcomes. Phenotypic changes in proliferation, migration, invasion, and differentiation will directly validate the functional relevance of the targeted GEPs and their constituent genes. The identification of specific driver genes within these programs will confirm their oncogenic roles and highlight potential therapeutic targets. Ultimately, the validation of these GEPs as functional entities will provide robust biomarkers for diagnosis, prognosis, and prediction of therapeutic response in pediatric cancers. This mechanistic understanding is crucial for the development of novel, targeted therapies that exploit the unique biological dependencies of pediatric tumors.

**Potential Challenges**

Several challenges may arise during these validation efforts. Inter-patient variability in PDX models can complicate interpretation and necessitate a large cohort of models. Technical limitations of spatial transcriptomics, such as resolution and throughput, may impact the ability to resolve fine-grained cellular interactions. Furthermore, off-target effects of CRISPR/Cas9 or RNAi perturbations could confound results, requiring careful validation of specificity. The complexity of GEPs, involving multiple co-regulated genes, means that targeting a single gene may not fully recapitulate the program's overall effect, necessitating combinatorial approaches.

**B. Immune Compartment**

**Scientific Rationale and Novelty**

Immune GEPs are hypothesized to reflect specific immune cell populations and their dynamic functional states within the tumor microenvironment (TME). Understanding these programs is critical for deciphering immune evasion mechanisms and for developing effective immunotherapies in pediatric cancers. The novelty lies in functionally validating the roles of these immune GEPs in shaping the anti-tumor immune response or promoting tumor progression, particularly in the context of specific pediatric tumor types.

**Experimental Approach**

To characterize immune cell subsets, their activation states, and their functional interactions within the TME, the following strategies are proposed:

1. **Immune Profiling**:
   - **Design**: Patient tumor samples and PDX models will be subjected to comprehensive immune profiling. Technologies such as multi-color flow cytometry and mass cytometry (CyTOF) will be employed to identify and quantify various immune cell subsets (e.g., T cells, B cells, NK cells, macrophages, dendritic cells, neutrophils). These techniques allow for deep phenotyping based on surface and intracellular markers. Re-analysis of scRNA-seq data, potentially with higher sequencing depth for immune cells, will further refine the characterization of immune cell states and their associated GEPs.
   - **Link to GEPs**: This approach will directly link specific immune GEPs (e.g., Program 1: Macrophage/Myeloid Activation; Program 9: B Cell-Like; Program 17: Macrophage/Osteoclast Differentiation; Program 19: T/B Cell Progenitor Immunity; Program 22: Humoral Immune Response; Program 26: Platelet/Megakaryocyte-Like; Program 27: Mast Cell Activation; Program 38: Neutrophil Granule/Myeloid Activation; Program 44: Plasmacytoid Dendritic Cell-Like; Program 57: T Cell Receptor Signaling; Program 67: Neutrophil Activation; Program 68: Dendritic Cell-Like; Program 69: Macrophage/Monocyte-Like; Program 70: Hypoxia/Acute Phase Response; Program 77: NK Cell Cytotoxicity; Program 87: Cytotoxic T Cell and NK Cell Effector; Program 108: T Cell Activation; Program 119: Interferon-Gamma Response; Program 121: B Cell and Plasma Cell Immune Response; Program 122: Inflammatory Cytokine and Chemokine Signaling; Program 129: Interferon Response; Program 134: Myeloid Cell/Neutrophil Innate Immunity) to their corresponding cell types and activation states within the tumor microenvironment.^1^
2. **Cytokine/Chemokine Analyses**:
   - **Design**: Quantification of secreted inflammatory mediators will be performed using multiplex ELISA, Luminex assays, or targeted proteomics on tumor lysates, conditioned media from co-cultures, or biofluids from PDX models. This will provide a functional readout of the inflammatory responses associated with specific immune GEPs.
   - **Link to GEPs**: This is particularly relevant for GEPs like Program 122 ("Non-malignant Inflammatory Cytokine and Chemokine Signaling"), which is characterized by key inflammatory cytokines and chemokines.^1^
3. **Cell-Cell Interaction Assays**:
   - **Design**: Co-culture experiments involving isolated immune cells and tumor or stromal cells will be established to investigate direct cellular interactions. Live-cell imaging will be employed to monitor dynamic processes such as immune cell migration, tumor cell killing, or immune synapse formation. Functional assays for T cell activation, cytokine production, and cytotoxic activity will be performed.
4. **Technology Integration for Immune Compartment**:
   - **Spatial Transcriptomics**: This technology is crucial for understanding the spatial distribution of immune cells and their GEPs relative to tumor cells and stromal elements. It will allow for the identification of immune "hotspots" or "coldspots" and the precise localization of immune-suppressive or immune-activating niches within the tumor.
   - **Patient-Derived Xenografts (PDX) with Humanized Immune Systems**: To accurately model human immune responses *in vivo*, humanized PDX models will be utilized. These models involve engrafting human hematopoietic stem cells into immunodeficient mice, allowing for the reconstitution of a functional human immune system that can then interact with the human tumor cells. This approach enables the study of human immune cell infiltration, activation, and anti-tumor efficacy in a physiologically relevant context.

**Expected Outcomes and Their Implications**

The proposed experiments are expected to provide a detailed characterization of the immune infiltrates in pediatric cancers, including the identification of specific immune cell subsets and their functional states. This will lead to the discovery of novel immune checkpoints or immune-modulatory pathways that contribute to tumor immune evasion. Understanding critical immune-malignant crosstalk will inform strategies to reprogram the TME for enhanced anti-tumor immunity. These findings have significant implications for developing new immunotherapeutic strategies, identifying predictive biomarkers for immunotherapy response, and potentially repurposing existing immune-modulating agents for pediatric cancer treatment.

**Potential Challenges**

Maintaining the viability and functional integrity of human immune cells in humanized PDX models can be challenging. The complex data analysis required for integrating multi-omics datasets from spatial transcriptomics and CyTOF necessitates advanced computational biology expertise. Ethical considerations surrounding the use of humanized models and human patient samples must also be carefully navigated. Furthermore, the inherent plasticity of immune cells means their functional state can be highly context-dependent, requiring careful experimental design to capture relevant biological insights.

**C. Stromal Compartment**

**Scientific Rationale and Novelty**

Stromal GEPs are hypothesized to reflect distinct stromal cell populations (e.g., fibroblasts, endothelial cells, mesenchymal stem cells) and their multifaceted roles in extracellular matrix (ECM) remodeling, angiogenesis, and providing supportive signals for tumor growth and metastasis. The novelty lies in functionally validating the specific pro-tumorigenic or anti-tumorigenic contributions of these stromal programs and identifying the key molecular players that mediate these effects within the pediatric tumor microenvironment.

**Experimental Approach**

To investigate the roles of stromal cells and their associated GEPs, the following experimental approaches are proposed:

1. **Fibroblast Characterization**:
   - **Design**: Isolation and characterization of cancer-associated fibroblasts (CAFs) from patient tumors and PDX models will be performed. This involves identifying specific CAF markers (e.g., α-SMA, FAP, PDGFRα/β) and assessing their functional properties *in vitro*, such as their ability to promote tumor cell proliferation, migration, or drug resistance.
   - **Link to GEPs**: This is directly relevant for programs like Program 2 ("Malignant OS - Bone Morphogenesis and Extracellular Matrix"), which is specific to Osteosarcoma and involves ECM components.^1^
2. **Extracellular Matrix (ECM) Analyses**:
   - **Design**: The composition and physical properties of the ECM will be assessed in tumor tissues. This includes immunohistochemistry for key ECM proteins (e.g., collagens, fibronectin, laminin), proteomics to identify the full ECM proteome, and biomechanical assays (e.g., atomic force microscopy, rheology) to measure tissue stiffness. These analyses will correlate with the activity of stromal GEPs involved in ECM remodeling.
   - **Link to GEPs**: Relevant genes in GEPs, such as collagens (e.g., COL11A1, COL9A1, COL27A1 in Program 120) ^1^, will be specifically investigated.
3. **Angiogenesis Assays**:
   - **Design**: *In vitro* assays, such as endothelial cell tube formation assays and spheroid sprouting assays, will be used to evaluate the pro-angiogenic potential of conditioned media from stromal cells or tumor cells with perturbed stromal GEPs. *In vivo* angiogenesis assays (e.g., Matrigel plug assay, corneal micropocket assay) will further confirm these findings.
4. **Technology Integration for Stromal Compartment**:
   - **Spatial Transcriptomics**: This technology is essential for mapping stromal GEPs within the tumor architecture, allowing for the precise localization of different stromal cell populations and their proximity to malignant and immune cells. This will reveal how stromal programs are spatially organized and how they contribute to distinct tumor microenvironments.
   - **Patient-Derived Xenografts (PDX) with Stromal Co-engraftment**: To model the complex tumor-stroma interactions *in vivo*, PDX models will be established with co-engraftment of human patient-derived stromal cells (e.g., CAFs, mesenchymal stem cells) alongside tumor cells. This approach enables the study of human stromal cell contributions to tumor growth, angiogenesis, and response to therapy in a more physiologically relevant setting.

**Expected Outcomes and Their Implications**

These experiments are expected to identify specific pro-tumorigenic or anti-tumorigenic stromal functions, leading to the validation of novel stromal targets for therapeutic intervention. Understanding the intricate tumor-stroma crosstalk will illuminate how the microenvironment supports tumor progression and metastasis. These findings have significant implications for developing therapies that target the supportive stromal niche, potentially sensitizing tumors to conventional treatments or preventing relapse. Furthermore, stromal GEPs may serve as novel biomarkers for disease progression or therapeutic response.

**Potential Challenges**

The isolation and successful long-term culture of primary stromal cells from patient samples can be technically challenging. The inherent heterogeneity of stromal cell populations within tumors complicates their functional characterization. Analyzing the complex composition and biomechanical properties of the ECM requires specialized expertise and equipment. Ethical considerations related to the use of human patient-derived stromal cells and their co-engraftment in animal models must be carefully addressed.

**V. Disease-Specific Experimental Plans**

This section outlines tailored experimental plans for various pediatric cancers, leveraging the identified GEPs and cutting-edge technologies, ordered by their prevalence in the dataset.

**A. Neuroblastoma (NB) - Prevalence: 29.380%**

**Why Neuroblastoma is of Interest**

Neuroblastoma (NB) is a highly aggressive pediatric cancer originating from neural crest cells, often presenting with complex genetic alterations and diverse clinical outcomes. The pan-cancer pediatric cohort scRNA-seq data has yielded a remarkable number of "Neurodevelopmental" GEPs highly specific to Neuroblastoma.^1^ These include Program 7 ("Neuronal Proliferation and Cholinergic Signaling"), Program 12 ("Schwann Cell-Like"), Program 15 ("Neuronal Synaptic and Calcium Signaling"), Program 33 ("Core Oncogenic/Proliferative - chr2p24 amplification"), Program 35 ("Neuronal Adhesion and Signaling"), Program 40 ("Neuroendocrine and Neuronal Differentiation"), Program 54 ("Neuronal Ion Homeostasis and Signaling"), Program 58 ("Mitochondrial Respiration and Axonal Transport"), Program 107 ("Synaptic Vesicle Dynamics and Neuronal Excitability"), Program 109 ("Synaptic Plasticity and Neuropeptide Signaling"), Program 110 ("Axonogenesis and Cholinergic Receptor"), and Program 111 ("Neuronal Axon Guidance").^1^ The presence of known oncogenic drivers like

*MYCN* and *ALK* within several of these GEPs further underscores their clinical relevance.^1^ Additionally, Neuroblastoma exhibits significant immune infiltration, as evidenced by numerous "Immune and Inflammatory Response" GEPs (e.g., Program 9, 17, 19, 22, 26, 27, 38, 44, 57, 67, 68, 69, 70, 77, 87, 108, 119, 121, 129, 134), highlighting the complex interplay between tumor cells and the immune microenvironment in this disease.^1^

**Unique Biological Questions Addressed**

The identified GEPs allow for addressing several unique biological questions in Neuroblastoma:

- How do specific neuronal GEPs, such as those involved in cholinergic signaling (Program 7) or calcium signaling (Program 15), contribute to Neuroblastoma aggressiveness and therapy resistance? ^1^
- What is the precise functional role of the "B cell-like" (Program 9) or "platelet/megakaryocyte-like" (Program 26) programs in Neuroblastoma malignancy? Are these truly tumor-intrinsic programs reflecting aberrant differentiation or lineage plasticity of Neuroblastoma cells, or do they represent the infiltration of non-malignant immune/hematopoietic cells that contribute to tumor progression? This distinction is critical for therapeutic targeting. ^1^
- How does the observed "Coordinated Malignant Neuroblastoma Identity," where multiple malignant neuronal GEPs (e.g., Program 15 and 110, Rho = 0.862; Program 33 and 111, Rho = 0.855; Program 54 and 109, Rho = 0.862) are highly co-expressed, function as a unified oncogenic network? Can targeting a master regulator within this network disrupt multiple facets of Neuroblastoma biology? ^1^

**Specific Experimental Plan for Neuroblastoma**

1. **Focus on Malignant Neuroblastoma GEPs (e.g., Program 7, 15, 33, 107, 109, 110, 111)**:
   - **Technology**: Spatial Transcriptomics and High-resolution Imaging.
   - **Design**: Neuroblastoma patient samples and established PDX models will be analyzed using high-resolution spatial transcriptomics. The aim is to precisely map the localization and co-expression patterns of highly correlated malignant Neuroblastoma GEPs (e.g., Program 15 and 110, Rho = 0.862; Program 33 and 111, Rho = 0.855; Program 54 and 109, Rho = 0.862) within the tumor architecture.^1^ This will involve correlating GEP activity with known histological features, regions of

*MYCN* amplification (for Program 7 and 33), and the expression of specific neuronal differentiation markers. High-resolution imaging techniques will complement spatial transcriptomics to visualize cellular morphology and spatial relationships.

- **Expected Outcomes**: Spatial validation of GEP co-expression patterns, identification of distinct malignant niches within the tumor, and visual confirmation of the spatial relationship between MYCN-driven proliferation and neuronal differentiation programs. This will provide a critical understanding of how these programs are organized *in situ*.

2. **Focus on Immune-Malignant Interactions (e.g., Program 87 & Malignant NB GEPs)**:
   - **Technology**: Humanized PDX models integrated with Multi-omics (scRNA-seq, CyTOF, Spatial Transcriptomics).
   - **Design**: Neuroblastoma tumor cells will be co-engrafted with specific human immune cell subsets (e.g., cytotoxic T cells and NK cells, which are highly active in Program 87) into immunodeficient mice with reconstituted human immune systems.^1^ Tumor growth and immune cell infiltration will be meticulously tracked. Spatial transcriptomics will then be applied to the resulting tumors to analyze the precise distribution of immune cells and their GEP activities in relation to the malignant Neuroblastoma GEPs. This will be complemented by scRNA-seq and CyTOF to deeply phenotype immune cells.
   - **Expected Outcomes**: A mechanistic understanding of immune evasion mechanisms in Neuroblastoma, identification of critical immune-malignant crosstalk pathways, and validation of potential immune-modulatory therapeutic targets that could enhance anti-tumor immunity.

**Potential Implications for Diagnostic, Prognostic, or Therapeutic Advancements**

These investigations hold significant potential for advancing Neuroblastoma management. The validated GEPs could serve as novel diagnostic biomarkers, allowing for more precise classification of tumor subtypes. Their activity might also correlate with clinical outcomes, providing improved prognostic markers. Most importantly, identifying and understanding the functional dependencies of these malignant neuronal GEPs and their interactions with the immune microenvironment will pave the way for developing novel therapeutic strategies, including targeted therapies that disrupt oncogenic neuronal pathways or immunotherapies that overcome immune suppression in Neuroblastoma. The clear distinction between tumor-intrinsic and microenvironmental GEPs will guide the development of more precise and effective treatments.

**B. Embryonal Rhabdomyosarcoma (ERMS) - Prevalence: 17.027%**

**Why Embryonal Rhabdomyosarcoma is of Interest**

Embryonal Rhabdomyosarcoma (ERMS) is a common soft tissue sarcoma in children, characterized by its resemblance to developing skeletal muscle. While the dataset highlights several immune-related GEPs enriched in ERMS, such as Program 1 ("Non-malignant Macrophage / Myeloid Activation" - ERMS: 0.8331), Program 108 ("Non-malignant T Cell Activation" - ERMS: 0.127), Program 119 ("Non-malignant Interferon-Gamma Response" - ERMS: 0.24), Program 121 ("Non-malignant B Cell and Plasma Cell Immune Response" - ERMS: 0.2006), Program 122 ("Non-malignant Inflammatory Cytokine and Chemokine Signaling" - ERMS: 0.1845), and Program 129 ("Non-malignant Interferon Response" - ERMS: 0.33) ^1^, there is a notable absence of ERMS-specific malignant GEPs in the provided annotation. This suggests that the primary insights from this dataset for ERMS may lie in understanding its immune microenvironment rather than distinct malignant cell programs.

**Unique Biological Questions Addressed**

Given the prevalence of immune GEPs in ERMS, unique biological questions include:

- How do the prominent macrophage/myeloid activation (Program 1) and T cell activation (Program 108) programs influence ERMS progression and response to therapy? ^1^
- What is the functional consequence of the strong interferon responses (Program 119, 129) and inflammatory cytokine signaling (Program 122) within the ERMS tumor microenvironment? Do these responses contribute to anti-tumor immunity or promote tumor growth and immune evasion? ^1^
- Can modulating these immune GEPs, particularly macrophage and T cell activities, be a viable therapeutic strategy for ERMS?

**Specific Experimental Plan for Embryonal Rhabdomyosarcoma**

1. **Characterization of Myeloid and T Cell Infiltrates**:
   - **Technology**: Spatial Transcriptomics, Multi-color Flow Cytometry/CyTOF, and Humanized PDX models.
   - **Design**: ERMS patient samples and humanized PDX models will be used to comprehensively profile immune cell infiltration. Spatial transcriptomics will map the precise localization of cells expressing Program 1 (macrophages/myeloid cells) and Program 108 (T cells) within the ERMS tumor architecture, correlating their presence with tumor cell proliferation and necrosis. Flow cytometry or CyTOF will quantify the activation states and phenotypes of these immune cells.
   - **Expected Outcomes**: Identification of specific immune cell subsets that are highly active in ERMS, understanding their spatial distribution relative to tumor cells, and insights into whether these immune infiltrates are pro-tumorigenic or anti-tumorigenic.
2. **Functional Impact of Inflammatory and Interferon Responses**:
   - **Technology**: *In vitro* co-culture systems, CRISPR/Cas9, and PDX models.
   - **Design**: ERMS cell lines will be co-cultured with immune cells (e.g., macrophages, T cells) under conditions that mimic the inflammatory and interferon responses indicated by GEPs 119, 122, and 129. Key genes from these GEPs (e.g., CXCL9, CXCL10, IL36B) will be perturbed in immune or stromal cells using CRISPR/Cas9 or shRNA. The impact on ERMS cell proliferation, migration, and immune cell-mediated killing will be assessed.
   - **Expected Outcomes**: Elucidation of the functional roles of specific inflammatory and interferon pathways in ERMS, identifying whether they contribute to tumor control or immune evasion, and pinpointing potential targets for immune modulation.

**Potential Implications for Diagnostic, Prognostic, or Therapeutic Advancements**

Understanding the immune landscape of ERMS through these GEPs could lead to novel immunotherapeutic strategies. Identifying specific immune cell subsets or inflammatory pathways that are either suppressed or aberrantly activated in ERMS could inform the development of immune checkpoint inhibitors, adoptive cell therapies, or cytokine-based treatments. These GEPs may also serve as prognostic biomarkers, predicting patient response to conventional therapies or identifying patients who might benefit from immune-modulating interventions.

**C. Osteosarcoma (OS) - Prevalence: 15.356%**

**Why Osteosarcoma is of Interest**

Osteosarcoma (OS) is the most common primary malignant bone tumor in children and adolescents, characterized by its aggressive nature and high metastatic potential. A key GEP identified in the pan-cancer cohort is Program 2 ("Malignant OS - Bone Morphogenesis and Extracellular Matrix"), which is highly specific to Osteosarcoma.^1^ This GEP's annotation points to its direct involvement in bone formation and ECM remodeling, processes central to Osteosarcoma pathogenesis. Furthermore, Osteosarcoma is also associated with "Immune and Inflammatory Response" GEPs, such as Program 122 ("Non-malignant Inflammatory Cytokine and Chemokine Signaling"), which shows the highest proportion of activating cells from Osteosarcoma (0.265), indicating a significant inflammatory component within its microenvironment.^1^

**Unique Biological Questions Addressed**

The identified GEPs allow for addressing several unique biological questions in Osteosarcoma:

- How does Program 2 specifically drive aberrant bone formation and extracellular matrix remodeling, which are hallmarks of Osteosarcoma progression and metastasis? ^1^
- What is the precise role of the stromal compartment, particularly as characterized by Program 2, in promoting Osteosarcoma growth, invasion, and drug resistance? ^1^
- How does the "Malignant OS - Bone Morphogenesis and Extracellular Matrix" GEP interact with the inflammatory programs prevalent in the Osteosarcoma microenvironment, such as Program 122 ("Inflammatory Cytokine and Chemokine Signaling")? Does this crosstalk contribute to tumor progression or immune evasion? ^1^

**Specific Experimental Plan for Osteosarcoma**

1. **Focus on Malignant OS GEP (Program 2)**:
   - **Technology**: PDX models and Spatial Transcriptomics.
   - **Design**: Established Osteosarcoma PDX models will be utilized. Key genes within Program 2 (e.g., those involved in bone morphogenesis and ECM remodeling, as indicated in its description) will be perturbed using CRISPR/Cas9 in Osteosarcoma cell lines. These perturbed cell lines will then be engrafted into mice to monitor primary tumor growth, metastatic burden, and the extent of aberrant bone formation or destruction. Spatial transcriptomics will be performed on the resulting PDX tumors to visualize the activity of Program 2 and its impact on the surrounding tumor microenvironment, including the host bone.
   - **Expected Outcomes**: Functional validation of Program 2's critical role in Osteosarcoma pathogenesis, identification of specific molecular drivers of aberrant bone remodeling, and the discovery of potential therapeutic targets that could inhibit tumor growth and metastasis by modulating this program.
2. **Focus on Stromal-Malignant Interactions (Program 2 & general stromal GEPs)**:
   - **Technology**: Spatial Transcriptomics and 3D Bioprinting/Organoids.
   - **Design**: Osteosarcoma patient samples and PDX models will be analyzed using spatial transcriptomics to comprehensively map the spatial relationship between Program 2 activity in malignant cells and the activity of other relevant stromal GEPs. Complementary to *in vivo* models, 3D bioprinting or organoid technologies will be employed to create *in vitro* Osteosarcoma models that incorporate both tumor cells and patient-derived stromal cells. These advanced models will recapitulate the complex bone microenvironment, enabling detailed studies of Program 2 interactions with stromal components and their functional consequences.
   - **Expected Outcomes**: A deeper understanding of the contribution of the stromal compartment to Osteosarcoma progression, identification of novel targets for disrupting pro-tumorigenic tumor-stroma crosstalk, and insights into how the physical and molecular properties of the ECM influence tumor behavior.

**Potential Implications for Diagnostic, Prognostic, or Therapeutic Advancements**

These investigations are poised to significantly impact Osteosarcoma management. The validated functional roles of Program 2 could lead to the development of highly specific targeted therapies for Osteosarcoma, moving beyond conventional chemotherapy. The GEPs could also serve as novel diagnostic markers for aggressive forms of Osteosarcoma or as prognostic indicators to predict disease recurrence or metastatic potential. Furthermore, understanding the intricate mechanisms by which Osteosarcoma cells interact with and manipulate their bone microenvironment will provide crucial insights into its metastatic processes, potentially leading to strategies that prevent or treat metastatic disease.

**D. Retinoblastoma (RB) - Prevalence: 7.558%**

**Why Retinoblastoma is of Interest**

Retinoblastoma (RB) is the most common intraocular malignancy in childhood. The dataset identifies two highly specific malignant GEPs: Program 39 ("Malignant RB - Retinal Glia and Lipid Metabolism") and Program 88 ("Malignant RB - Retinal Neurodevelopment and Oxidative Phosphorylation").^1^ These GEPs highlight unique aspects of RB biology, including its origin from retinal cells, aberrant neurodevelopmental pathways, and distinct metabolic dependencies, particularly oxidative phosphorylation.

**Unique Biological Questions Addressed**

The identified GEPs allow for addressing unique biological questions in Retinoblastoma:

- How do the retinal glial and neurodevelopmental programs (Program 39, 88) contribute to the malignant phenotype and growth of Retinoblastoma cells? ^1^
- What is the functional significance of the prominent lipid metabolism (Program 39) and oxidative phosphorylation (Program 88) in Retinoblastoma? Can these metabolic vulnerabilities be therapeutically exploited? ^1^
- Are there specific co-expression patterns between these two RB-specific GEPs that indicate a coordinated oncogenic program?

**Specific Experimental Plan for Retinoblastoma**

1. **Functional Validation of Retinal Neurodevelopment and Metabolism**:
   - **Technology**: *In vitro* 3D retinal organoids/spheroids, CRISPR/Cas9, and Metabolomics.
   - **Design**: Retinoblastoma cell lines or patient-derived cells will be cultured in 3D retinal organoid models to better mimic the tissue architecture. Key genes from Program 39 (e.g., *FABP7*, *APOE*) and Program 88 (e.g., *SLC25A4*, *UQCRHL*) will be perturbed using CRISPR/Cas9. Functional assays will assess cell proliferation, survival, and differentiation. Metabolomic profiling (e.g., mass spectrometry) will be performed to quantify changes in lipid profiles and OXPHOS intermediates following GEP perturbation.
   - **Expected Outcomes**: Confirmation of the role of specific genes and pathways in driving malignant retinal neurodevelopment and identifying metabolic vulnerabilities that can be targeted.
2. **Spatial Mapping of Metabolic Heterogeneity**:
   - **Technology**: Spatial Transcriptomics and Imaging Mass Cytometry (IMC).
   - **Design**: Retinoblastoma patient samples will be analyzed using spatial transcriptomics to map the distribution of cells activating Program 39 and Program 88 within the tumor. IMC will be used to visualize the spatial distribution of key metabolic enzymes and lipids, correlating them with GEP activity and cellular morphology.
   - **Expected Outcomes**: Understanding the spatial organization of metabolic programs within RB tumors and identifying distinct metabolic niches that could influence drug delivery or resistance.

**Potential Implications for Diagnostic, Prognostic, or Therapeutic Advancements**

Validation of these GEPs could lead to novel targeted therapies for Retinoblastoma that exploit its unique metabolic dependencies, potentially offering less toxic alternatives to conventional treatments. The GEPs could also serve as biomarkers for disease progression, response to therapy, or even predict the likelihood of metastasis.

**E. Alveolar Rhabdomyosarcoma (ARMS) - Prevalence: 7.269%**

**Why Alveolar Rhabdomyosarcoma is of Interest**

Alveolar Rhabdomyosarcoma (ARMS) is an aggressive subtype of rhabdomyosarcoma with a poor prognosis. The dataset identifies Program 118 ("Malignant ARMS - Myogenesis / Neural Development") as highly specific to ARMS, uniquely highlighting both myogenic and neural developmental features.^1^ This mixed-lineage characteristic is a critical biological question. Additionally, ARMS shows a significant presence of Program 27 ("Non-malignant Mast Cell Activation / Allergic Response" - ARMS: 0.5556), suggesting an important role for mast cells in its microenvironment.^1^

**Unique Biological Questions Addressed**

The identified GEPs allow for addressing unique biological questions in ARMS:

- How does the co-activation of myogenic and neural developmental programs (Program 118) contribute to the aggressive phenotype and developmental plasticity of ARMS cells? ^1^
- What are the master regulators that drive this mixed-lineage differentiation in ARMS, and can targeting them induce terminal differentiation or inhibit tumor growth?
- What is the functional role of mast cell infiltration (Program 27) in the ARMS tumor microenvironment? Do mast cells promote tumor growth, angiogenesis, or immune suppression? ^1^

**Specific Experimental Plan for Alveolar Rhabdomyosarcoma**

1. **Dissecting Myogenic and Neural Plasticity**:
   - **Technology**: Single-cell multi-omics (scRNA-seq + scATAC-seq), CRISPR/Cas9, and PDX models.
   - **Design**: ARMS cell lines and patient-derived cells will be subjected to scRNA-seq and scATAC-seq to identify individual cells co-expressing myogenic (e.g., *TNNT2*) and neural (e.g., *NOS1*, *DCX*) markers from Program 118, and to map their epigenetic landscape. CRISPR/Cas9 will be used to perturb key transcription factors or signaling pathways hypothesized to drive this mixed-lineage state. The impact on differentiation, proliferation, and tumor formation in PDX models will be assessed.
   - **Expected Outcomes**: Identification of regulatory mechanisms underlying ARMS[ developmental plasticity and potential targets to induce differentiation or block aberrant development.
2. **Role of Mast Cells in ARMS Microenvironment**:
   - **Technology**: Spatial Transcriptomics, Co-culture assays, and Humanized PDX models.
   - **Design**: Spatial transcriptomics will map the spatial relationship between ARMS tumor cells and mast cells (identified by Program 27 markers like *TPSAB1*, *CPA3*) within patient samples and humanized PDX models. Co-culture experiments with ARMS cells and mast cells will investigate direct interactions, assessing mast cell-mediated effects on tumor cell growth, migration, and angiogenesis.
   - **Expected Outcomes**: Understanding the contribution of mast cells to ARMS progression and identifying potential targets for modulating mast cell activity to enhance anti-tumor responses.

**Potential Implications for Diagnostic, Prognostic, or Therapeutic Advancements**

Understanding the developmental plasticity of ARMS could lead to novel differentiation therapies. Targeting mast cell infiltration or their pro-tumorigenic functions could represent a new immunotherapeutic approach for ARMS. These GEPs could also serve as biomarkers for ARMS subtypes with specific differentiation states or microenvironmental characteristics.

**F. Ewing Sarcoma (EWS) - Prevalence: 3.904%**

**Why Ewing Sarcoma is of Interest**

Ewing Sarcoma (EWS) is an aggressive bone and soft tissue sarcoma, primarily driven by *EWSR1-ETS* fusions. Despite its mesenchymal origin, the dataset reveals a striking and consistent activation of "Neurodevelopmental" GEPs in EWS, including Program 16 ("Malignant EWS - Neuronal Differentiation and Synaptic Modulation"), Program 92 ("Malignant EWS - Cell Adhesion/Synapse Assembly"), Program 97 ("Malignant EWS - Neural Developmental Regulation"), and Program 112 ("Malignant EWS - Neuronal Action Potential and Calcium Signaling").^1^ This convergent neuronal phenotype is a significant and non-obvious finding.

**Unique Biological Questions Addressed**

The identified GEPs allow for addressing unique biological questions in EWS:

- How do these distinct neuronal GEPs (Program 16, 92, 97, 112) collectively contribute to the malignant phenotype of Ewing Sarcoma, given its mesenchymal origin? Is this a form of aberrant differentiation or lineage plasticity? ^1^
- Can targeting specific neuronal signaling pathways (e.g., calcium signaling from Program 112) or cell adhesion molecules (from Program 92) inhibit EWS growth and metastasis? ^1^
- What are the co-regulation mechanisms between these highly correlated EWS neuronal GEPs (e.g., Program 16 and 112, Rho = 0.890; Program 16 and 97, Rho = 0.5439)? Do they share common upstream regulators? ^1^

**Specific Experimental Plan for Ewing Sarcoma**

1. **Functional Validation of Neuronal Phenotypes**:
   - **Technology**: CRISPR/Cas9, Functional Assays (2D/3D), and PDX models.
   - **Design**: EWS cell lines will be used to perturb key genes identified in the neuronal GEPs (e.g., *SOX6*, *PAX7*, *PAX3*, *NKX2-2* from Program 16; *CACNA1I*, *RYR2* from Program 112; *HMCN1*, *CDH8* from Program 92). Functional assays will assess changes in proliferation, migration, invasion, and neuronal differentiation markers (e.g., neurite outgrowth, expression of neuronal proteins). Perturbed cells will be engrafted into PDX models to assess *in vivo* tumor growth and metastatic potential.
   - **Expected Outcomes**: Confirmation of the functional contribution of neuronal programs to EWS malignancy and identification of specific neuronal targets for therapeutic intervention.
2. **Investigating Cross-GEP Co-regulation**:
   - **Technology**: ChIP-seq, ATAC-seq, and High-throughput CRISPR screens.
   - **Design**: For highly correlated EWS GEPs (e.g., Program 16 and 112), ChIP-seq and ATAC-seq will identify shared transcription factor binding sites and open chromatin regions. High-throughput CRISPR screens will then target candidate master regulators (e.g., transcription factors, epigenetic modifiers) to identify those that simultaneously affect the expression of multiple EWS neuronal GEPs.
   - **Expected Outcomes**: Elucidation of the shared regulatory mechanisms that drive the coordinated activation of neuronal programs in EWS, potentially revealing novel pan-neuronal targets.

**Potential Implications for Diagnostic, Prognostic, or Therapeutic Advancements**

The validation of these neuronal GEPs could lead to a paradigm shift in understanding EWS biology, moving beyond its mesenchymal classification. This could open avenues for repurposing neuro-active drugs or developing novel therapies that target these aberrant neuronal pathways. The identified master regulators could serve as novel diagnostic or prognostic biomarkers and as targets for more effective and less toxic treatments.

**G. Malignant Rhabdoid Tumor (MRT) - Prevalence: 3.043%**

**Why Malignant Rhabdoid Tumor is of Interest**

Malignant Rhabdoid Tumor (MRT) is a highly aggressive pediatric cancer, often characterized by *SMARCB1* (INI1) inactivation. Program 37 ("Malignant MRT - Developmental / Cell Adhesion") is highly specific to MRT, indicating a strong association with developmental processes and cell adhesion.^1^ This aligns with the understanding of MRT as a tumor arising from primitive cells with disrupted differentiation.

**Unique Biological Questions Addressed**

The identified GEP allows for addressing unique biological questions in MRT:

- How does the "Developmental / Cell Adhesion" program (Program 37) contribute to the aggressive growth and metastatic potential of MRT cells? ^1^
- What are the specific roles of genes like *OPCML*, *CNTN4*, and *PTCH1* within this GEP in driving MRT pathogenesis? ^1^
- Can targeting components of this developmental program induce differentiation or sensitize MRT cells to existing therapies?

**Specific Experimental Plan for Malignant Rhabdoid Tumor**

1. **Functional Characterization of Developmental and Adhesion Genes**:
   - **Technology**: CRISPR/Cas9, 3D Spheroid/Organoid models, and PDX models.
   - **Design**: MRT cell lines will be used to perturb key genes from Program 37 (e.g., *OPCML*, *CNTN4*, *PTCH1*, *FGFR2*). Functional assays will assess the impact on cell proliferation, self-renewal (in 3D spheroid/organoid models), migration, and invasion. Perturbed cells will be engrafted into PDX models to evaluate *in vivo* tumor growth and metastasis.
   - **Expected Outcomes**: Elucidation of the functional roles of specific developmental and cell adhesion genes in MRT malignancy and identification of potential therapeutic targets.

**Potential Implications for Diagnostic, Prognostic, or Therapeutic Advancements**

Understanding the developmental programs active in MRT could lead to novel differentiation therapies or strategies to restore normal developmental cues. The identified GEP components could serve as biomarkers for MRT diagnosis, prognosis, or response to targeted therapies.

**H. Desmoplastic Small Round Cell Tumor (DSRCT) - Prevalence: 2.215%**

**Why Desmoplastic Small Round Cell Tumor is of Interest**

Desmoplastic Small Round Cell Tumor (DSRCT) is a rare but highly aggressive sarcoma, often characterized by the *EWSR1-WT1* fusion. The dataset identifies three highly specific malignant GEPs for DSRCT: Program 62 ("Malignant DSRCT - Neuronal and Astrocytic Development"), Program 66 ("Malignant DSRCT - Neural Crest / Sympathoadrenal"), and Program 82 ("Malignant DSRCT- Neuronal Adhesion / Synapse Organization").^1^ These GEPs strongly suggest a neural crest or neuroectodermal origin and highlight neuronal differentiation, astrocytic features, and synaptic organization.

**Unique Biological Questions Addressed**

The identified GEPs allow for addressing unique biological questions in DSRCT:

- How do the neural crest and sympathoadrenal developmental programs (Program 66) contribute to the aggressive and metastatic nature of DSRCT? ^1^
- What is the functional significance of the neuronal and astrocytic development (Program 62) and neuronal adhesion/synapse organization (Program 82) in DSRCT malignancy? Do these programs represent therapeutic vulnerabilities? ^1^
- Can targeting specific genes like *NTRK3* (from Program 82) or pathways like calcium signaling (from Program 62) inhibit DSRCT growth? ^1^

**Specific Experimental Plan for Desmoplastic Small Round Cell Tumor**

1. **Investigating Neural Crest and Neuronal Differentiation**:
   - **Technology**: CRISPR/Cas9, Functional Assays, and PDX models.
   - **Design**: DSRCT cell lines will be used to perturb key genes from Programs 62, 66, and 82 (e.g., *GLI2*, *NTRK3*, *ADGRB1*). Functional assays will assess changes in proliferation, migration, invasion, and expression of neural crest or neuronal differentiation markers. Perturbed cells will be engrafted into PDX models to evaluate *in vivo* tumor growth and metastatic potential.
   - **Expected Outcomes**: Elucidation of the functional roles of neural crest and neuronal developmental programs in DSRCT malignancy and identification of specific targets for therapeutic intervention.
2. **Spatial Mapping of Neural Features**:
   - **Technology**: Spatial Transcriptomics.
   - **Design**: DSRCT patient samples will be analyzed using spatial transcriptomics to map the precise localization of cells activating Programs 62, 66, and 82 within the tumor. This will help determine if these neuronal features are uniformly expressed or localized to specific tumor niches.
   - **Expected Outcomes**: Understanding the spatial heterogeneity of neural differentiation in DSRCT and its potential implications for tumor behavior and treatment response.

**Potential Implications for Diagnostic, Prognostic, or Therapeutic Advancements**

The validation of these GEPs could provide novel insights into the developmental origins of DSRCT, leading to new diagnostic markers. Targeting specific neural developmental pathways or genes like *NTRK3* could offer new therapeutic avenues for this aggressive cancer.

**I. Wilms Tumor (WT) - Prevalence: 3.482%**

**Why Wilms Tumor is of Interest**

Wilms Tumor (WT) is the most common kidney cancer in children, believed to arise from aberrant kidney development. The dataset identifies Program 59 ("Malignant WT - Neuroepithelial Development") as highly specific to Wilms Tumor, indicating a neuroepithelial developmental component.^1^ This is interesting given its kidney origin and suggests developmental plasticity.

**Unique Biological Questions Addressed**

The identified GEP allows for addressing unique biological questions in Wilms Tumor:

- How does the "Neuroepithelial Development" program (Program 59) contribute to the malignant phenotype and developmental characteristics of Wilms Tumor cells? ^1^
- What is the functional significance of genes like *RFX4*, *SOX2-OT*, and *NPAS3* in driving this neuroepithelial program in WT? ^1^
- Can targeting this neuroepithelial developmental pathway induce differentiation or inhibit tumor growth in WT?

**Specific Experimental Plan for Wilms Tumor**

1. **Functional Validation of Neuroepithelial Development**:
   - **Technology**: CRISPR/Cas9, 3D Organoid models (e.g., kidney organoids), and PDX models.
   - **Design**: Wilms Tumor cell lines or patient-derived cells will be used to perturb key genes from Program 59 (e.g., *RFX4*, *SOX2-OT*). These cells will be incorporated into 3D kidney organoid models to assess their ability to form organized structures and differentiate. Functional assays will evaluate proliferation, self-renewal, and differentiation markers. Perturbed cells will be engrafted into PDX models to assess *in vivo* tumor growth.
   - **Expected Outcomes**: Confirmation of the role of neuroepithelial developmental genes in WT malignancy and identification of potential targets for differentiation therapy.

**Potential Implications for Diagnostic, Prognostic, or Therapeutic Advancements**

Understanding the neuroepithelial developmental program in WT could lead to novel differentiation-inducing therapies. The GEP could also serve as a biomarker for specific WT subtypes or for predicting response to developmental pathway inhibitors.

**J. Undifferentiated Round Cell Sarcoma (URCS) - Prevalence: 1.309%**

**Why Undifferentiated Round Cell Sarcoma is of Interest**

Undifferentiated Round Cell Sarcoma (URCS) is a rare and aggressive sarcoma. The dataset identifies Program 74 ("Malignant URCS - Mesenchymal and Neural Development") as highly specific to URCS, uniquely integrating both kidney epithelial and neural developmental characteristics.^1^ This mixed-lineage phenotype is a significant biological finding.

**Unique Biological Questions Addressed**

The identified GEP allows for addressing unique biological questions in URCS:

- How does the co-activation of kidney epithelial and neural developmental programs (Program 74) contribute to the malignant phenotype and developmental plasticity of URCS cells? ^1^
- What are the master regulators that drive this mixed-lineage differentiation in URCS, and can targeting them induce terminal differentiation or inhibit tumor growth?
- What is the functional significance of genes like *HMGA2*, *ETV1*, and *CLDN16* in this context? ^1^

**Specific Experimental Plan for Undifferentiated Round Cell Sarcoma**

1. **Dissecting Mixed Mesenchymal and Neural Plasticity**:
   - **Technology**: Single-cell multi-omics (scRNA-seq + scATAC-seq), CRISPR/Cas9, and PDX models.
   - **Design**: URCS cell lines and patient-derived cells will be subjected to scRNA-seq and scATAC-seq to identify individual cells co-expressing kidney epithelial and neural markers from Program 74, and to map their epigenetic landscape. CRISPR/Cas9 will be used to perturb key transcription factors or signaling pathways hypothesized to drive this mixed-lineage state. The impact on differentiation, proliferation, and tumor formation in PDX models will be assessed.
   - **Expected Outcomes**: Identification of regulatory mechanisms underlying URCS developmental plasticity and potential targets to induce differentiation or block aberrant development.

**Potential Implications for Diagnostic, Prognostic, or Therapeutic Advancements**

Understanding the developmental plasticity of URCS could lead to novel differentiation therapies. The identified GEP components could serve as biomarkers for URCS subtypes with specific differentiation states or for predicting response to developmental pathway inhibitors.

**K. High-Grade Sarcoma (HGS) - Prevalence: 1.158%**

**Why High-Grade Sarcoma is of Interest**

High-Grade Sarcoma (HGS) represents a diverse group of aggressive mesenchymal tumors. The dataset identifies two malignant GEPs with strong enrichment in HGS: Program 14 ("Malignant HGS - Neuronal Signaling and Development") and Program 79 ("Malignant Neural / Renal Developmental - HGS and MRT Enriched").^1^ These GEPs suggest an unexpected neuronal component in a subset of HGS, similar to observations in Ewing Sarcoma.

**Unique Biological Questions Addressed**

The identified GEPs allow for addressing unique biological questions in HGS:

- How do the neuronal signaling and developmental programs (Program 14, 79) contribute to the aggressive phenotype of HGS, given its mesenchymal origin? Is this a shared mechanism with other sarcomas exhibiting neuronal features? ^1^
- What is the functional significance of genes like *GRM7*, *DAB1*, and *PDE10A* (Program 14) or *NPHS1*, *ETV4*, and *NEUROG2* (Program 79) in driving HGS malignancy? ^1^
- Can targeting these neuronal pathways offer new therapeutic avenues for HGS?

**Specific Experimental Plan for High-Grade Sarcoma**

1. **Functional Validation of Neuronal Signaling**:
   - **Technology**: CRISPR/Cas9, Functional Assays, and PDX models.
   - **Design**: HGS cell lines that activate Program 14 and/or 79 will be used to perturb key genes involved in neuronal signaling and development. Functional assays will assess changes in proliferation, migration, and invasion. The impact on tumor growth and metastasis will be evaluated in PDX models.
   - **Expected Outcomes**: Confirmation of the functional contribution of neuronal programs to HGS malignancy and identification of specific neuronal targets for therapeutic intervention.
2. **Comparative Analysis with other Neuronal Sarcomas**:
   - **Technology**: Comparative Spatial Transcriptomics.
   - **Design**: Spatial transcriptomics data from HGS tumors activating neuronal GEPs will be compared with similar data from Ewing Sarcoma and DSRCT to identify common spatial patterns of neuronal differentiation and their relationship to tumor architecture and microenvironment.
   - **Expected Outcomes**: Identification of shared vulnerabilities across sarcomas exhibiting neuronal phenotypes, potentially leading to pan-sarcoma therapeutic strategies.

**Potential Implications for Diagnostic, Prognostic, or Therapeutic Advancements**

The identification of neuronal GEPs in HGS could lead to a reclassification of certain HGS subtypes and the development of targeted therapies that exploit these unexpected neuronal dependencies. These GEPs could also serve as diagnostic or prognostic biomarkers.

**L. Embryonal Sarcoma (EMBSARC) - Prevalence: 0.315%**

**Why Embryonal Sarcoma is of Interest**

Embryonal Sarcoma (EMBSARC) is a rare mesenchymal tumor. The dataset identifies Program 85 ("Malignant EMBSARC - Neural / Developmental") as predominantly associated with EMBSARC, highlighting neural adhesion and developmental features.^1^ This suggests a developmental origin or plasticity in these tumors.

**Unique Biological Questions Addressed**

The identified GEP allows for addressing unique biological questions in EMBSARC:

- How does the "Neural / Developmental" program (Program 85) contribute to the malignant phenotype and growth of EMBSARC cells? ^1^
- What are the specific roles of genes like *CDH9*, *CNTN5*, and *MYRF* in driving this developmental program in EMBSARC? ^1^
- Can targeting this neural developmental pathway induce differentiation or inhibit tumor growth in EMBSARC?

**Specific Experimental Plan for Embryonal Sarcoma**

1. **Functional Validation of Neural/Developmental Program**:
   - **Technology**: CRISPR/Cas9, 3D Spheroid/Organoid models, and PDX models.
   - **Design**: EMBSARC cell lines or patient-derived cells will be used to perturb key genes from Program 85. Functional assays will assess the impact on cell proliferation, self-renewal, migration, and differentiation. Perturbed cells will be engrafted into PDX models to evaluate *in vivo* tumor growth.
   - **Expected Outcomes**: Confirmation of the role of neural and developmental genes in EMBSARC malignancy and identification of potential therapeutic targets.

**Potential Implications for Diagnostic, Prognostic, or Therapeutic Advancements**

Understanding the neural/developmental program in EMBSARC could lead to novel differentiation-inducing therapies. The GEP could also serve as a biomarker for EMBSARC diagnosis or for predicting response to developmental pathway inhibitors.

**M. Other Pediatric Cancers**

For the remaining pediatric cancer types in the dataset (ACT, CCSK, IFS, CCSST, SCS, SRMS, GCT, GIST, EPSARC, SS), no specific GEPs were identified in the provided annotation table.^1^ Therefore, detailed experimental validation plans cannot be proposed based on the current research material. Future research would require additional GEP identification and annotation for these tumor types to develop targeted experimental strategies.

**VI. Cross-GEP Interactions**

The co-expression matrix provides compelling evidence of functional relationships between GEPs, extending beyond individual programs to suggest coordinated biological processes. Investigating these cross-GEP interactions is crucial for a holistic understanding of pediatric cancer biology.

**A. Co-regulation Mechanisms**

**Scientific Rationale and Novelty**

Highly co-expressed GEPs are hypothesized to be regulated by shared underlying mechanisms, such as common transcription factors, epigenetic modifiers, or central signaling pathways. Identifying these shared regulatory nodes represents a novel approach to understanding the coordinated activation of multiple biological programs in cancer. This moves beyond single-gene or single-pathway targeting to address broader regulatory vulnerabilities.

**Experimental Approach**

To elucidate the co-regulation mechanisms underlying highly correlated GEPs, an integrated multi-omics approach is proposed:

1. **Integrated Multi-omics (ChIP-seq, ATAC-seq, Proteomics)**:
   - **Design**: For highly correlated GEP pairs (e.g., Program 15 & 110, Program 33 & 111, Program 54 & 109 in Neuroblastoma; Program 16 & 112 in Ewing Sarcoma) ^1^, chromatin immunoprecipitation sequencing (ChIP-seq) will be performed to identify common transcription factor binding sites (TFBS) across the co-expressed GEPs. Assay for Transposase-Accessible Chromatin using sequencing (ATAC-seq) will map regions of open chromatin, indicating active regulatory elements. Proteomic analysis (e.g., quantitative mass spectrometry) will identify co-expressed proteins and activated signaling pathways that may serve as upstream regulators.
   - **CRISPRi/a Screens**: High-throughput CRISPR interference (CRISPRi) or CRISPR activation (CRISPRa) screens will be conducted in relevant pediatric cancer cell lines. These screens will target predicted master regulators (e.g., transcription factors, kinases) identified from the multi-omics data. The impact of perturbing these regulators on the expression levels of multiple co-expressed GEPs will be simultaneously assessed using multiplexed reporter assays or targeted RNA sequencing.

**Expected Outcomes and Their Implications**

These experiments are expected to identify master regulators that govern the coordinated activity of multiple GEPs. This will lead to the elucidation of shared regulatory networks that underpin complex malignant phenotypes. The discovery of such central nodes offers novel therapeutic targets, as modulating a single master regulator could effectively disrupt multiple interdependent oncogenic programs, potentially leading to more potent and durable anti-cancer effects.

**B. Synergistic or Antagonistic Roles in Disease Contexts**

**Scientific Rationale and Novelty**

Beyond shared regulation, co-expressed GEPs may exert synergistic or antagonistic functional roles in disease progression. Understanding these complex functional interactions is critical for designing effective combinatorial therapies and anticipating potential compensatory mechanisms that lead to drug resistance. The novelty lies in experimentally dissecting these functional relationships in a disease-relevant context.

**Experimental Approach**

To explore the synergistic or antagonistic roles of co-expressed GEPs, the following approaches are proposed:

1. **Dual Perturbation Studies (CRISPR/Cas9 + Pharmacologic Inhibitors)**:
   - **Design**: Highly correlated GEP pairs (e.g., Program 14 & 117, Program 65 & 128, Program 16 & 112) will be selected for investigation.^1^ In relevant pediatric cancer cell lines, genes from one GEP will be perturbed using CRISPR/Cas9 (knockout or inhibition), and simultaneously, pharmacologic inhibitors targeting key pathways or genes within the co-expressed GEP will be applied. The combinatorial effects on critical tumor phenotypes (e.g., proliferation, survival, migration, differentiation) will be rigorously assessed. This approach will reveal synergistic vulnerabilities or antagonistic interactions that could inform novel drug combinations.
2. **Spatial Multi-omics (e.g., MERFISH, Vizgen MERSCOPE)**:
   - **Design**: To understand the spatial context of GEP interactions, high-resolution spatial multi-omics technologies will be employed on pediatric patient tumor samples. Techniques such as Multiplexed Error-Robust Fluorescence *in situ* Hybridization (MERFISH) or Vizgen MERSCOPE allow for single-cell resolution mapping of hundreds to thousands of RNA transcripts within intact tissue sections. This will enable the identification of individual cells co-activating multiple GEPs and their precise spatial relationships with other cell types and tissue structures.
   - **Expected Outcomes**: These spatial analyses will reveal whether co-expressed GEPs are active within the same cells or in spatially distinct but interacting cell populations. This will provide unprecedented insights into the cellular and architectural context of GEP synergy or antagonism, informing how these interactions contribute to tumor heterogeneity and function *in situ*.

**Expected Outcomes and Their Implications**

These experiments are expected to identify novel synergistic drug combinations that could overcome therapeutic resistance. They will also shed light on compensatory mechanisms, where the inhibition of one GEP might lead to the upregulation of a co-expressed GEP, thereby informing strategies to prevent resistance. The spatial mapping of GEP interactions will provide a critical understanding of how these programs contribute to specific tumor microenvironments and ultimately, to disease progression. This knowledge is paramount for developing more effective and spatially informed therapeutic strategies.

**VII. Conclusions and Future Directions**

The comprehensive analysis of 134 GEPs derived from single-cell RNA-seq data of a pan-cancer pediatric cohort offers a profound opportunity to advance the understanding and treatment of childhood cancers. The proposed experimental validation plans, meticulously designed for malignant, immune, and stromal compartments, and tailored to specific diseases like Neuroblastoma, Osteosarcoma, Retinoblastoma, Alveolar Rhabdomyosarcoma, Ewing Sarcoma, Malignant Rhabdoid Tumor, Desmoplastic Small Round Cell Tumor, Wilms Tumor, Undifferentiated Round Cell Sarcoma, Embryonal Sarcoma, and High-Grade Sarcoma, represent a critical step in translating computational discoveries into biological and clinical realities.

The identification of highly correlated GEPs, particularly within malignant Neuroblastoma programs, suggests a tightly integrated oncogenic network rather than isolated biological processes. This implies that targeting master regulators within such networks could yield more effective and durable therapeutic responses. The recurrent observation of neuronal phenotypes in various sarcomas, despite their diverse mesenchymal origins, points to a convergent oncogenic differentiation, potentially revealing pan-sarcoma vulnerabilities. The nuanced interpretation of "malignant" labels for certain immune/hematopoietic GEPs underscores the necessity of rigorous experimental validation to distinguish between tumor-intrinsic programs and infiltrating non-malignant cells, a distinction with direct implications for therapeutic strategy. Furthermore, the prominent role of oxidative phosphorylation in specific malignant neuronal tumors challenges a universal view of cancer metabolism, opening new avenues for metabolic targeting. Lastly, the presence of mixed-lineage and developmental plasticity GEPs highlights the inherent adaptability of pediatric tumors, suggesting that targeting aberrant developmental pathways could be a potent therapeutic approach.

The integration of cutting-edge technologies like spatial transcriptomics and patient-derived xenografts is central to these plans. Spatial transcriptomics will provide unprecedented insights into the *in situ* organization and interactions of GEPs within the complex tumor microenvironment, while PDX models will enable physiologically relevant *in vivo* validation of functional hypotheses and therapeutic efficacy.

The successful execution of these validation plans is anticipated to yield a wealth of knowledge that can be directly translated into clinical advancements. The validated GEPs and their key driver genes hold immense potential as novel diagnostic biomarkers for precise tumor classification, prognostic indicators for predicting disease course, and, most critically, as actionable therapeutic targets. By moving from correlative observations to mechanistic understanding, these efforts will pave the way for the development of more personalized, effective, and less toxic treatments for children battling cancer. Future directions will involve expanding these validation efforts to a broader spectrum of pediatric cancers, exploring the dynamic evolution of GEPs under therapeutic pressure, and ultimately, translating these laboratory findings into clinical trials to improve patient outcomes.

FOLLOW-UP:

Major findings out of the dataset so far (none of these are binding – right now, we’re just throwing things out there):

1. Neuroblastoma tumors consist of many intermixed GEPs spanning a variety of neuronal and neuroendocrine processes

- While this isn’t ground-breaking, it would be worth looking to see if any of these programs correlate with high-risk disease or poor prognosis.
- We could use the TARGET bulk RNA-seq dataset (which has both RNA-seq and survival data).
- We could apply RNA velocity to our neuroblastoma malignant data to estimate transitions between cell states
- Comparison of MYCN-specific program (#7), ALK-specific program (#15), and dual MYCN+ALK program (#33)
- How do specific neuronal GEPs, such as those involved in cholinergic signaling (Program 7) or calcium signaling (Program 15), contribute to Neuroblastoma aggressiveness and therapy resistance?
- We could overlap GEPs onto spatial omics datasets that we’ve already generated...

2. A cryptic neuronal program is active across a broad range of mesenchymal tumors, driven by transcriptional dysregulation.

- What are unifying features in the GEPs across the sarcoma neuronal programs? Can we find/predict a collection of drivers and/or TFs? This could potentially be validated in vitro.
- Can we apply spatial proteomics to a broad range of clinical samples? We’d need to make a TMA, but we’ve already validated a few neuronal marker antibodies with Jasmine and the CSO.
- Is this targetable? GEMINI suggests calcium channel modulators, but we’d have to look closely at these GEPS and see if there’s a reasonable argument there...

3. A unique mast cell program is present in fusion-positive ARMS.

- I double-checked – it's really there, and not specific to one sample. ARMS has a unique fibrotic structure, and I wonder if this ties into the NOS1 expression and tissue architecture of the disease.
- We could stain TMAs of RMS for mast cell markers.
- We could try to use bulk deconvolution of old RMS expression data to see if mast cell presence correlates with outcome
- I guess the question is – where do we go from there? I want to avoid getting committed to a complicated in vivo experiment mixing tumor cells and mast cells...

4. Metabolic reprogramming in embryonal tumors

- It appears that some tumors, especially neuroblastoma express a GEP that is enriched for mitochondrial and ox phos genes.
- Are these GEPs expressed in areas of hypoxia?

5. Tumors can “garble” GEPs to create hybrid cell states

- Are these truly new and hybrid states? Can we overlay these GEPs in reference atlas of normal development to show that they are separate in normal tissue but intermingled in cancers?

6. ERMS-specific questions:

- How do the prominent macrophage/myeloid activation (Program 1) and T cell activation (Program 108) programs influence ERMS progression and response to therapy? ^1^
- What is the functional consequence of the strong interferon responses (Program 119, 129) and inflammatory cytokine signaling (Program 122) within the ERMS tumor microenvironment? Do these responses contribute to anti-tumor immunity or promote tumor growth and immune evasion? ^1^
- Can modulating these immune GEPs, particularly macrophage and T cell activities, be a viable therapeutic strategy for ERMS?
